# Patient-matched analysis identifies deregulated networks in prostate cancer to guide personalized therapeutic intervention

**DOI:** 10.1101/695999

**Authors:** Akinchan Kumar, Alaa Badredine, Karim Azzag, Yasenya Kasikçi, Marie Laure Quintyn Ranty, Falek Zaidi, Nathalie Serret, Catherine Mazerolles, Bernard Malavaud, Marco Antonio Mendoza-Parra, Laurence Vandel, Hinrich Gronemeyer

## Abstract

Prostate cancer (PrCa) is the second most common malignancy in men^1^. More than 50% of advanced prostate cancers display the TMPRSS2-ERG fusion^2^. Despite extensive cancer genome/transcriptome^2–4^ and phosphoproteome^5^ data, little is known about the impact of mutations and altered transcription on regulatory networks in the PrCa of individual patients. Using patient-matched normal and tumor samples, we established somatic variations and differential transcriptome profiles of primary ERG-positive prostate cancers. Integration of protein-protein interaction and gene-regulatory network databases^6, 7^ defined highly diverse patient-specific network alterations. We found that different components of a given regulatory pathway were altered by novel and known mutations and/or aberrant gene expression, including deregulated ERG targets, such that different sets of pathways were altered in each individual PrCa. In a given PrCa, several deregulated pathways share common factors, predicting synergistic effects on cancer progression. Our integrated analysis provides a paradigm to identify key deregulated factors within regulatory networks to guide personalized therapies.

---

The mutational landscapes of primary and advanced/metastatic PrCa have been extensively analyzed^2, 3, 8, 9^, as has been the prevalence of the androgen-sensitive TMPRSS2 promoter fusion with ETS transcription factors^10^, which endows ETS with responsiveness to the androgen receptor (AR) that is frequently overexpressed in antiandrogen-resistant PrCa^11^. Recurrent mutations have been found in genes coding for factors regulating a plethora of pathways and key cellular functions, such as the androgen receptor signaling, PI3K/RAS/RAF/WNT pathways, and factors involved in DNA repair and chromatin methylation, or cell cycle control. One of the caveats in all these studies was that, with a few exceptions^9, 12^, information was generally compiled from large numbers of tumors from different patients. Thus, while enabling identification of predominant mutations, these studies did not reveal the spectrum of aberrations that existed in individual patients’ prostates at diagnosis. All these aberrations may affect different regulatory pathways and their added, possibly synergistic action may be critical for malignancy and tumor progression. Indeed, restoring a normal state will require the correction of a highly complex and dynamically regulated system of interactive multi-component networks which are deregulated in disease^13^. Towards this goal, the identification of aberrant networks and their inherent hierarchies is essential to design patient-selective therapeutic interventions through generic or key factor-specific modulation of the affected pathways.

We chose prostate cancer as a solid tumor paradigm to integrate patient-specific differential expressed (DEGs) and mutated genes, using information from protein-protein interaction and gene-regulatory network databases^6, 7^ (Fig. 1a) to generate patient-specific cancer-modified networks. Extensively characterized normal and tumor frozen punch biopsies from the same prostate were obtained from radical prostatectomy specimens of non-treated patients. 15 primary ERG-positive tumors (T) and matched normal tissue (N) were selected by expert pathologists on the basis that consecutive sections of the same biopsy differed only minimally in tumor cellularity (>80% tumor cells), while the sections of N biopsies from the same prostate had 0% tumor cells. With one exception of patient 14 (P14), the proportion of infiltrating lymphocytes relative to tumor cells was close to 0%, only occasionally rare scattered lymphocytes were observed in the stroma. The tumor sections of P14 showed up to 25% (area-based) mononuclear immune cells. Immunohistochemistry and RNA-seq confirmed ERG overexpression relative to the matched N samples and all samples revealed increased androgen receptor (AR) levels (Fig. 1b, c).

**Fig. 1:**
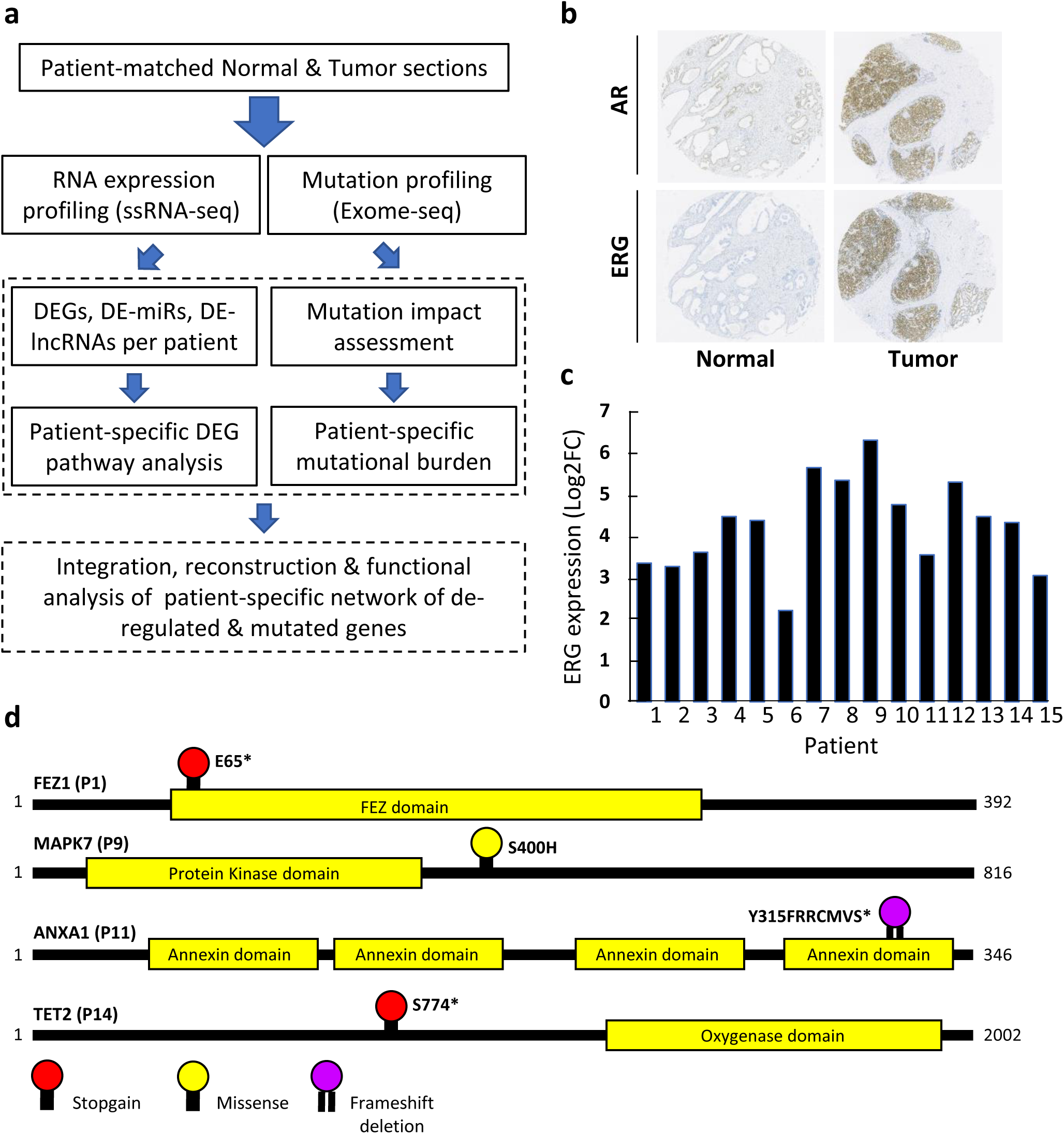
Analysis strategy and characterization of patient-matched samples. **a**, Sketch of the workflow of this study. **b**, Representative IHC images of cancer and corresponding matched normal samples from patient 15 stained with an anti-AR antibody (top panel) or an anti-ERG antibody (bottom panel). **c**, RNA-seq data of ERG expression in all the patients’ tumors relative to their matched normal. Differential ERG expression for all patient-matched duplicate samples was supported by q value < 10^−69^ using DEseq (see Methods for details). **d,** Schematic illustration of 4 novel mutations in P1, P9, P11 and P14, which are predicted to have a high or moderate impact.

Whole-exome sequencing (WES, Supplementary Table 1) and subsequent calling of variants by MuTect2 revealed between 49 and 114 mutations in each cancer relative to the corresponding normal prostate tissue; only mutations predicted to have high or moderate impact were considered subsequently (Supplementary File 1). Intriguingly, in addition to classical mutations, for example in *MYC*, *TP53, PTEN* or components of the PI3K and WNT pathways^2, 9, 14^, unreported patient-specific mutations were observed in all samples (Supplementary Table 2; for validations see Extended Data Fig. 1). In P1 three hitherto unreported somatic mutations affected the putative tumor suppressors *BANP*^15^, *FEZ1*^16, 17^ (Fig. 1d) and *TINAGL1*, which interferes with both integrin and EGFR signaling^18^. We also found novel mutations in *MAPK7* (R400H) in P9, Annexin A1 (*ANXA1*, frameshift deletion; P11) and a *TET2* mutation that truncates the protein and renders it non-functional (Fig. 1d; P14). These novel somatic mutations were seen only in single patients. However, the nature of the mutations, often truncating proteins of functional importance, is likely to have a significant impact in the individual case. Indeed, the ability of TINAGL1 to inhibit progression and metastasis of triple-negative breast cancer^18^, provides strong rational for such personalized genomic analysis. Our data underscores the recent notion that “significantly mutated genes” in PrCa may occur frequencies of only a few percent^19^.

Mutations in regulatory elements (e.g., enhancers) and factors (e.g., transcription factors, epigenetic modulators, enzymes) can affect a plethora of pathways. To integrate these effects in the network analysis, we performed duplicate high-throughput strand-specific paired-end total RNA sequencing after ribosomal RNA depletion from matched T and N biopsy sections. As expected, T *vs*. N analysis of the RNA-seq datasets identified tumor-specific differentially expressed genes (TS-DEGs; Supplementary File 2) with diverse functionalities, comprising (i) cancer-specific deregulated proto-oncogenes like c-*MYC* (all except P6, P11, P13) but also (ii) pleiotropic factors like the serine protease KLK4 (P4, P10, P14, P15), a regulator of AR and the PI3K/AKT/mTOR pathway^20^ and of protease-activated receptors^21^. Notably, deletion of *KLK4* impairs PrCa growth^20^. Moreover, (iii) epigenetic modifiers like *JMJD6* (P14), *KDM4B* (P5), *KDM6A* (P2, P8), *KDM6B* (P6, P9), *TET3* (P12, P14), *KAT2A* (P3), *KAT6A* (P2) or HDAC9 (P2-5, P7-11, P13-15) were differentially expressed in certain tumors. In addition to protein-coding genes, also the expression of (iv) certain regulatory RNAs was altered in tumors [micro-RNAs (miRs), as well as long non-coding RNA (lncRNAs); for annotated miRs and lncRNAs, see Supplementary Table 3]. Of note, the p53-inducible lncRNA NEAT, a promising therapeutic target whose ablation generates synthetic lethality with chemotherapy and p53 reactivation therapy^22, 23^, was over-expressed in 7/15 PrCa samples. A prominent ERG binding site in VCaP and in normal prostate epithelial RWPE-1 cells about 4.6kb upstream of the NEAT transcriptional start site may account for this deregulation (see Methods). The androgen-responsive lncRNA ARLNC1^24^ was up-regulated in 9/15 paired samples but down-regulated in P5. HOTTIP, a component of H3K4 methyltransferase complexes^25^ that can act as AR co-activator^26^ and was reported as negatively androgen-regulated lncRNA^24^ in prostate cancer cells, was downregulated in 9/15 PrCa samples. This included several, but not all of those with up-regulated ARLNC1. A similar divergence was seen with putative tumor suppressor and oncogenic miRNAs that are actively considered for clinical development^27^. For example, the RNA levels of tumor-suppressor miR34a were decreased in three samples (P10, P13 and P15) but increased in P4 and not affected in 11 other samples. MiR222, which displays targetable oncoMir characteristics in liver, pancreas and lung tumors^27, 28^, was unexpectedly down-regulated in 8/15 PrCa samples. Together, these vastly divergent genetic mutations and altered, often counter-intuitive gene expression patterns revealed the need to decipher for each individual patient the complexity of the deregulated systems to identify key targets in critical signaling pathways and/or key nodes in (sub)networks for concomitant intervention at several functionally different levels to generate synergistic effects.

As a first step towards the integration of the various deregulated functions within each tumor, we performed a patient-centered pathways enrichment analysis for TS-DEGs using Panther^29^ in the GenCodis3 environment^30^. While this analysis revealed several pathways commonly deregulated in PrCa of several patients - particularly cadherin, Wnt and integrin signaling - it also demonstrated that in each patient different sets of pathways were deregulated. Indeed, P5 and P6 had, respectively, the most and least severely affected PrCa in terms of numbers of deregulated pathways (Fig. 2; Extended Data Fig. 2 shows additional 52 patients from the TCGA repository). Moreover, different numbers and components of a commonly deregulated pathway were altered in different patients, yielding different p-values. As pointed out previously^14^, while genetic mutations of core Wnt pathway components are rare in PrCa, abnormal expression of β−catenin is frequent, suggesting that this deregulation occurs indirectly.

**Fig. 2:**
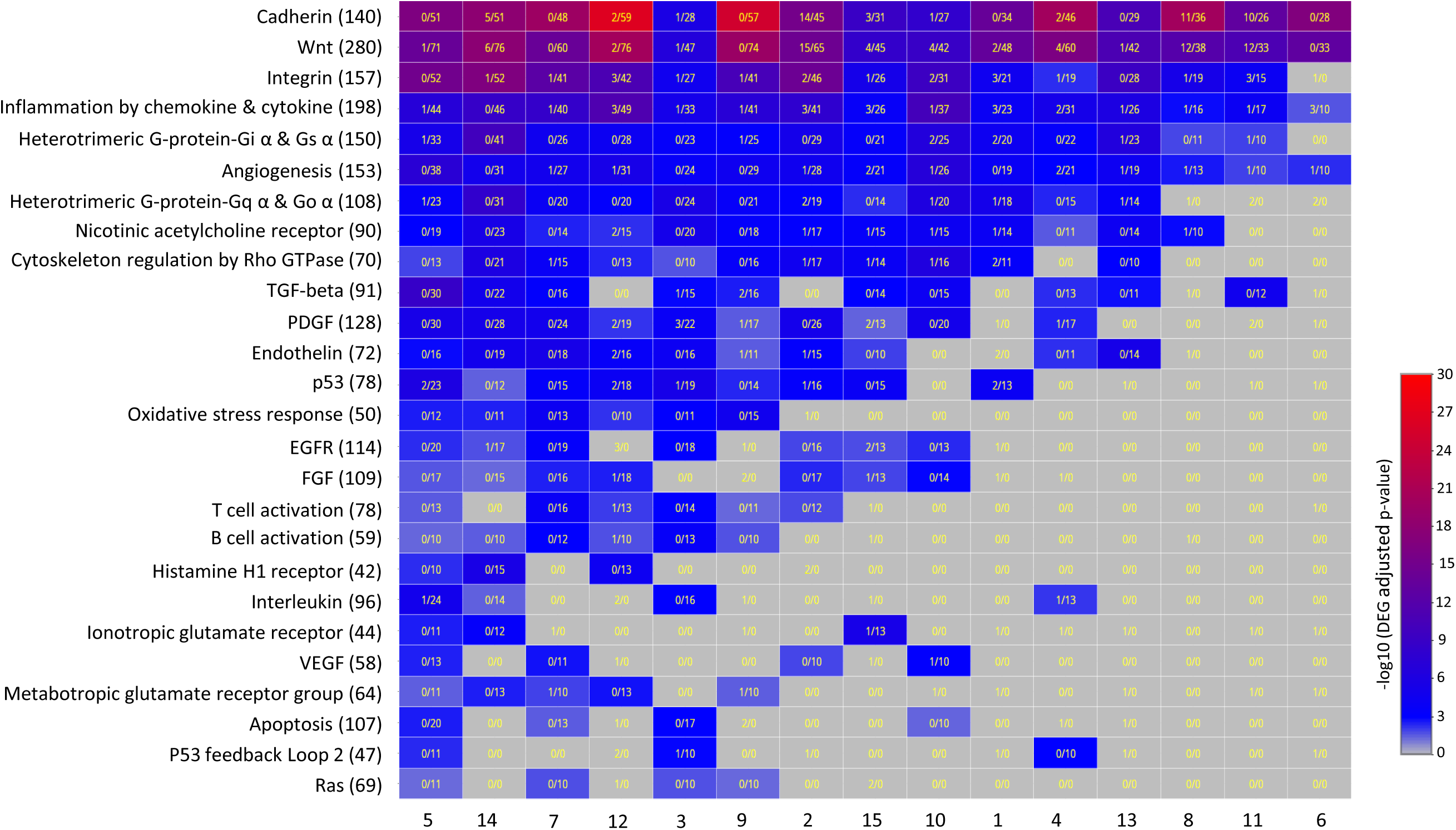
Divergence of pathways and severity of pathway alteration in individual prostate cancer patients. Pathways predicted by Panther to be significantly enriched in patient-specific DEGs are shown on the left of the table while patient identification numbers are given at the bottom. Panther-computed p-values for the deregulation of a given pathway are illustrated as blue-to-red color-coded rectangles; grey color indicated no significant alteration. Numbers in the rectangles give the mutated and deregulated genes for each pathway and patient (mutated/DEG; see Methods for details).

Genes never function in isolation but rather in a highly complex physiological context, which can be illustrated by their “communication” with other cellular components. To gain a more precise insight into the altered “communication” by patient-specific gene deregulations and mutations, we re-constructed “master networks” from all deregulated for each prostate cancer by integrating the connectivities provided by the STRING protein-protein and CellNet transcription factor-target gene interaction databases; in addition, we integrated all mutated genes and identified putative ERG and AR target genes by cognate binding sites in the vicinity of the transcriptional start site (TSS) [see Methods; Supplementary File 3 (cytoscape masterfile for each patient)]. Within these master networks, we studied first the components of the canonical and non-canonical Wnt pathways by merging all 183 deregulated/mutated genes of 15 patients (Extended Data Fig. 3a). Displaying the affected components in color in the context of the entire Wnt pathway connectivity revealed an unexpected heterogeneity (Fig. 3, Extended Data Fig. 3b-l). In P2 (Fig. 3a) an important signaling factor (phospholipase PLCB1) for the production of second messenger molecules (DAG, IP3) is mutated in the phospholipase domain (V571M) and the expression of multiple other master genes is deregulated, including *PPP3CA*, *GSK3B*, *MYC*, *TP53*, *HDAC1* in addition to several *WNT* and *Frizzled (FZD)* receptor genes. Notably, ChIP-seq data of TMPRSS2-ERG positive VCaP and normal RWPE1 prostate epithelial cells indicate that *WNT7B* and *HDAC1* are putative dual AR and ERG target genes, most likely affected by deregulated ERG and possibly, AR signaling (Extended Data Fig. 4a, b). Even more strikingly, the genes of several key signaling factors (PAK1, CREM) and of the epigenetic modulator SMARCD3 have apparently acquired ERG binding capability in their promoter regions during tumorigenesis (for *SMARCC1,* see Extended Data Fig. 4c), as it was reported for the ERG-mediated repression of checkpoint kinase 1^31^. In contrast, P6 showed a very small number of deregulated components of the core Wnt pathway (Fig. 3b), comprising three upregulated *FZD* receptors along with the cognate *WNT2B* ligand and *WNT2* which acquired ERG binding near the TSS in VCaP cells (Extended Data Fig. 4c, e). While such a scenario may be addressed with WNT inhibitor-based therapeutics, the diversity of deregulated Wnt networks in different patients may explain why the notion that targeting high Wnt-β-catenin signaling in cancer would be universally beneficial has been called into question^32^. P9 and P10 revealed two other scenarios of individual network alterations (Fig. 3c, d). Such patient-specific network alteration was also seen for less frequently affected signaling pathways. The PDGF and EGFR pathways were affected seriously in 10 and 7 patients, respectively (Fig. 4a, e; merged networks of alterations). However, the scenarios were completely different across individual patients (Fig. 4; Extended Data Fig. 5). Important changes were seen in P4 and P5 (Fig. 4b, c) but hardly any in P13 (Fig. 4d). The same was true for alterations of the EGFR pathway in P5 and P15 (Fig. 4f, g), while much less nodes were affected in P8 (Fig. 4h).

**Fig. 3:**
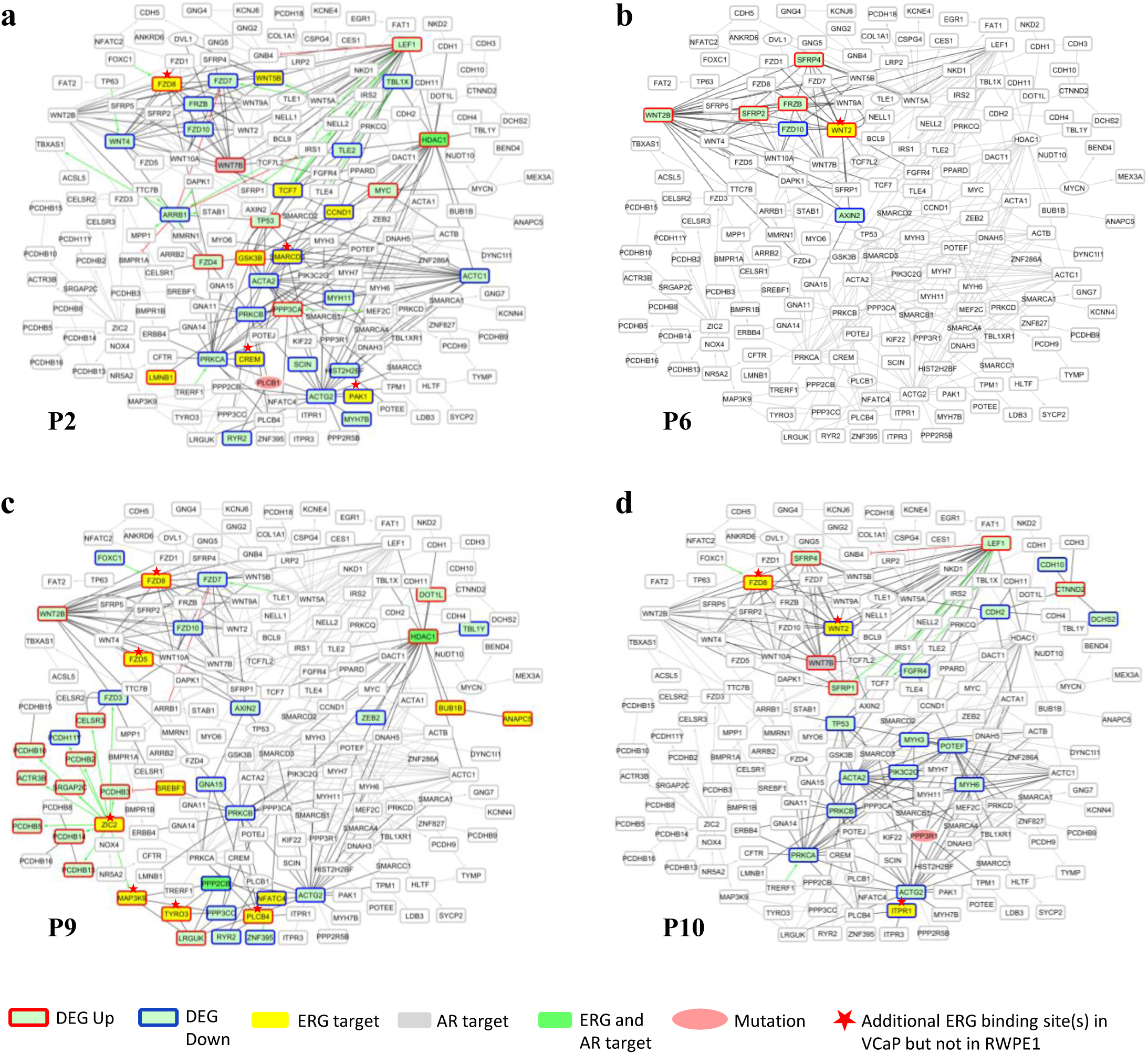
Patient-specific aberrations of the Wnt network are highly divergent. Global networks were established from differentially expressed genes (DEGs, >2-fold) in duplicate patient-matched tumor *vs.* normal samples by integrating the connectivities provided by STRING (protein-protein interaction database) and CellNet (transcription factor-target gene interactions); this integration yielded a ‘master network’ of DEGs for each patient, revealing the connectivities between deregulated genes. The DEG master network of each patient was complemented by the mutations of predicted high and moderate impact, and the components of the canonical and non-canonical Wnt pathways were extracted. DEGs and mutated genes are depicted in color for **a,** P2, **b,** P6, **c,** P9 and **d,** P10 in the background (grey nodes and connectivities) of all merged components of the Wnt pathways that are deregulated or mutated in all 15 patients (Suppl. Fig. 3a). The corresponding deregulated networks of the other patients are shown in Suppl. Fig. 3b-l). When known, connectivities are displayed as green (activation) or red (inhibition) lines; unknown connectivities and protein-protein interactions they are displayed as grey lines. DEG specifics and mutations are color-coded as describe below the figure.

**Fig. 4:**
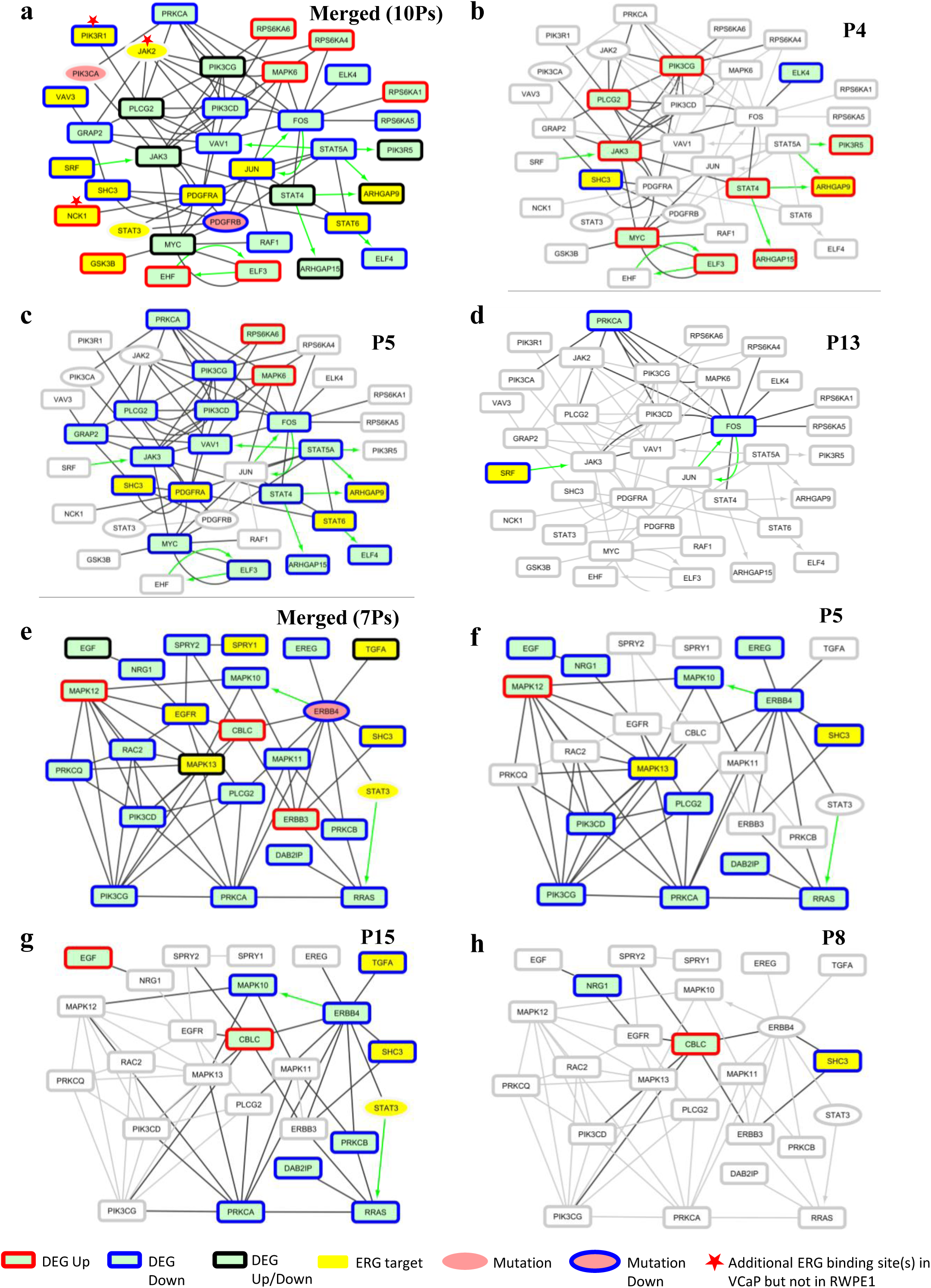
Divergent alterations of the PDGF and EGFR signaling networks in each patient. Ten patient exhibited serious aberrations in the PDGF and 7 patients in the EGFR signaling networks. **a,** Merged network of alterations (DEG, mutation) in the PDGF and **e,** EGFR networks. Using this merged network as background (grey nodes and connectivities) the aberrations in each individual patient are depicted in color. **b, c,** Patients with heavily (P4, P5) or **d,** minimally (P13) affected PDGF networks. **f, g,** Patients with heavily (P5, P15) or **h,** minimally (P8) affected EGFR networks. Color codes are displayed below the figure and in the legend to Fig. 3.

Finally, given that pathways do not act in isolation, we extracted the affected components of several pathways from the “master networks”. This analysis showed very clearly that, for P2 and P5 several genes of the Wnt, cadherin and integrin pathways, are shared between two or even three pathways (Fig. 5); the same was observed for other combinations of pathways (Extended Data Fig. 6). The functional consequence of deregulation/mutation of such genes is predicted to be serious and such nodes may comprise candidates for therapeutic targeting. It is worth pointing out that also genes at the nexus of several pathways diverged from one patient to another, as shown for P2 and P10 (Fig. 5a, b). Indeed, a hypothetical treatment of these two patients - assuming that drugs targeting key components would be available - will have to consider very different scenarios. In the PrCa of P2, common to two or all three pathways, there is a strong upregulation of the expression of several *WNT* and *FZD* genes, as well as *GSK3B* and *LEF1*. All these factors are potentially druggable and clinical trials are pursued at various levels. In addition, these genes are functionally connected with important other upregulated genes of the Wnt-pathway, such as *TP53*, *MYC*, *HDAC1* or *PPP3CA*. For P10, only two *FZD* genes are overexpressed in cancer and all deregulated *WNT* genes are less expressed than in the normal prostate tissue of this patient. Moreover, *HDAC1* is mutated and *MYC* is rather repressed. On the other hand, *RANBP2* is uniquely overexpressed in P10. Given its multi-functional role as E3-SUMO protein ligase, its overexpression may be an important component of the deregulated network.

**Fig. 5:**
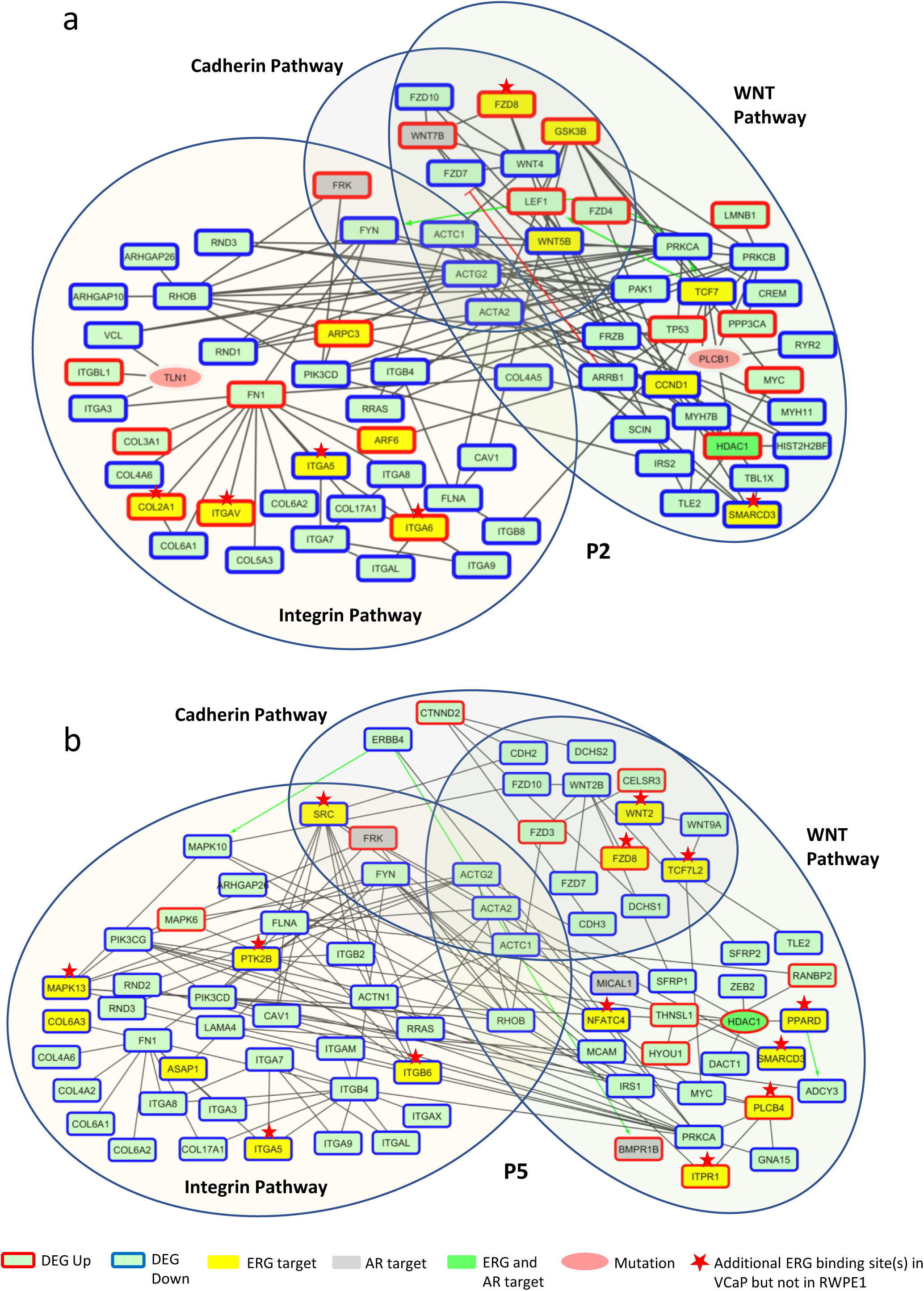
Affected signaling network crosstalk divergently between each other in each patient. **a,** Illustration of the merged networks of the affected genes from the Wnt, Cadherin and Integrin signaling pathways in the prostate of P2 revealing that several of the DEGs are common to different pathways. **b,** Illustration as in **a** but for P10. Color codes are displayed below the figure and in the legend to Fig. 3.

Taken together, the patient-centered network analysis we describe here for TMPRSS2-ERG positive primary prostate cancer reveals very divergent patient-specific deregulated and mutated genomic landscapes. This supports a rationale in which therapeutic options are considered in the context of a personalized integrative functional genomics analysis.

In the present case, the affected organs have been surgically removed and with one exception, all patients are still alive. However, we conducted this study as paradigm for other solid cancers where surgery is not possible and for monitoring the development of resistance during therapy. In addition, there is a growing importance of single cell functional genomics done with circulating tumor cells for diagnosis. Ultimately, additional dimensions, such as chromatin accessibility or RNA regulators such as the newly described circular RNAs^33, 34^, as well as metabolomic changes, may be integrated in this analysis to reveal what communication networks are at the origin, maintenance and progression of the disease and which regulatory circuits can be modulated for therapeutic purposes, including escape from resistance to therapy.

## Online Content

Any methods, additional references, Nature Research reporting summaries, source data, statements of code and data availability and associated accession codes are available at …

## Acknowledgments

We dedicate this work to the memory of Dr. Catherine Mazerolles who sadly died in 2015 when this study was gaining full speed. We thank all members of the Gronemeyer lab for discussions and suggestions. YK was supported by a fellowship of the EU Erasmus program. These studies were supported by funds from the Plan Cancer, AVIESAN-ITMO Cancer to HG and LV, the Ligue National Contre le Cancer (HG; Equipe Labellisée); and the Institut National du Cancer (INCa) to HG and LV. Support of the Agence Nationale de la Recherche (ANRT-07-PCVI-0031-01, ANR-10-LABX- 0030-INRT and ANR-10-IDEX-0002-02) is acknowledged.

## Author contributions

FZ and NS prepared the tissue sections for genomic, transcriptomic and TMA analyses; CM, ML-Q and NS performed the histopathological analyses of normal and tumor prostate samples. CM and BM supervised the clinical aspects of the study. KA performed all exome-seq and AK all transcriptome experiments. AK did together with AB all bioinformatics analyses under the guidance of HG and MAM-P. YK contributed to the experimental work and did all validation experiments for MuTect2-predicted mutations. HG and LV conceived and designed the study and wrote the manuscript with the help of AK, AB, MAM-P and BM.

## Additional information

**Extended data** is available for this paper at …

**Supplementary information** is available for this paper at …

**Reprints and permissions information** is available at …

**Correspondence and request for materials** should be addressed to H.G.

---

**Extended Data Fig. 1:**
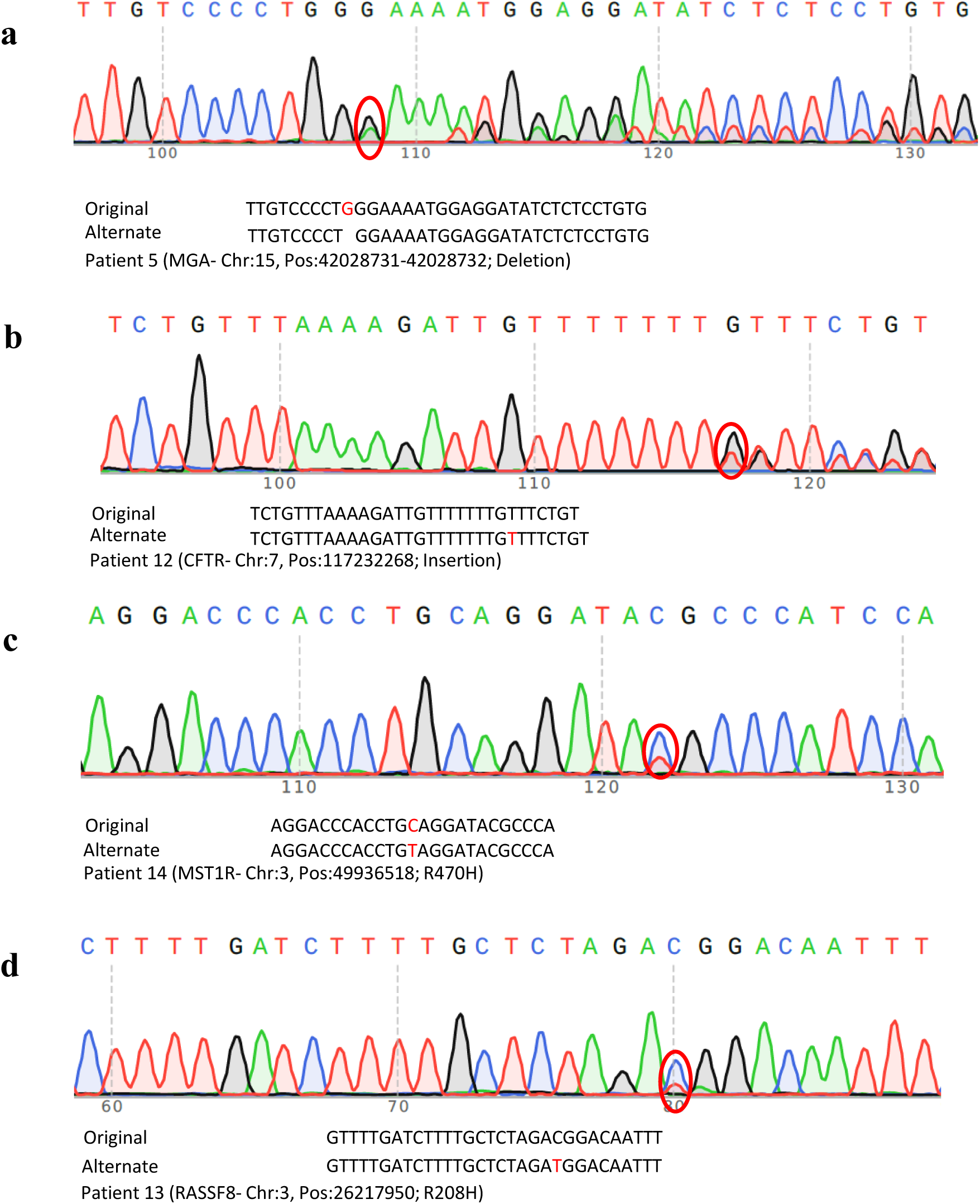
Validation of four selected different types of mutations from Exome-seq analysis using PCR coupled to Sanger sequencing. **a,** Single base deletion in *MGA* gene in patient 5. **b,** Single base insertion in *CFTR* in patient 12. **c,** Missense mutation in *MST1R* in patient 14 and **d,** Point mutation in *RASSF8* in patient 13. Red ovals highlight the regions of the mutations. Original (color-coded at the top) and mutated sequences are depicted below each sequence for comparison.

**Extended Data Fig. 2:**
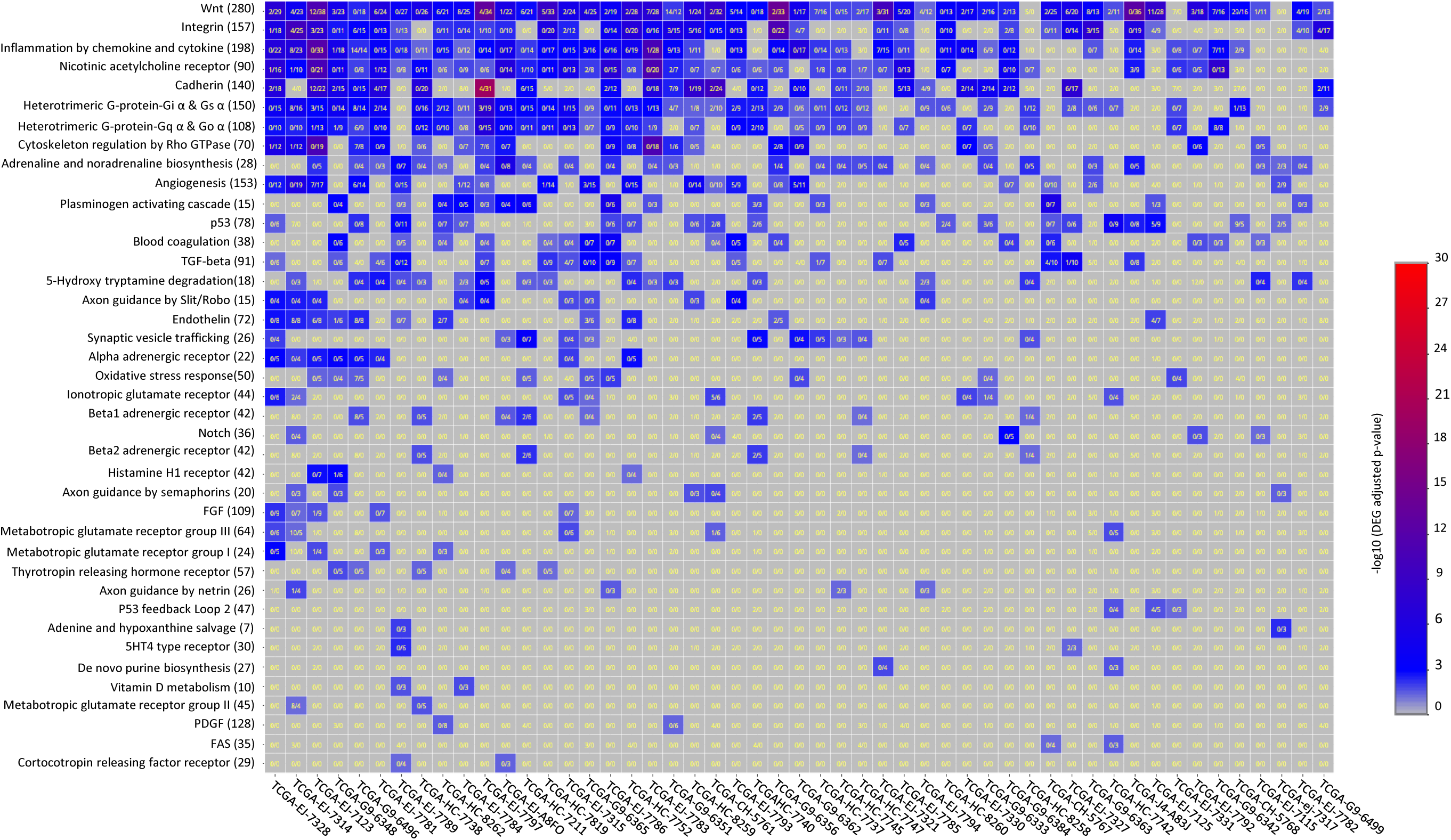
Heatmap showing enriched pathways using gene expression data from TCGA for prostate cancer. Note that we selected only datasets for which PrCa with patient-matched normal tissue were available. P-values corresponding to PANTHER-predicted alteration of pathway (left) are shown in color code (scale is shown on the right of the table). TCGA codes for patients are given at the bottom.

**Extended Data Fig. 3:**
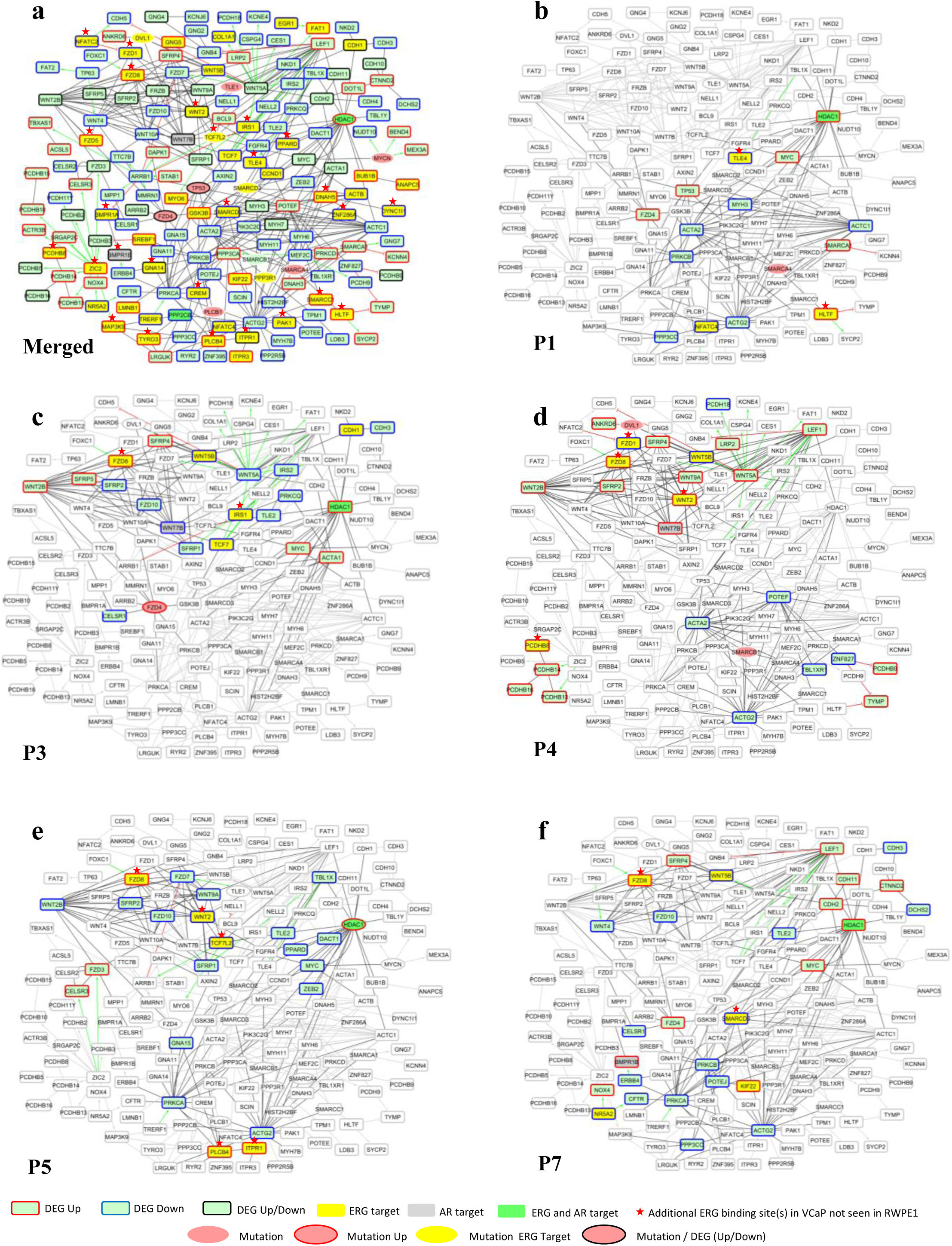
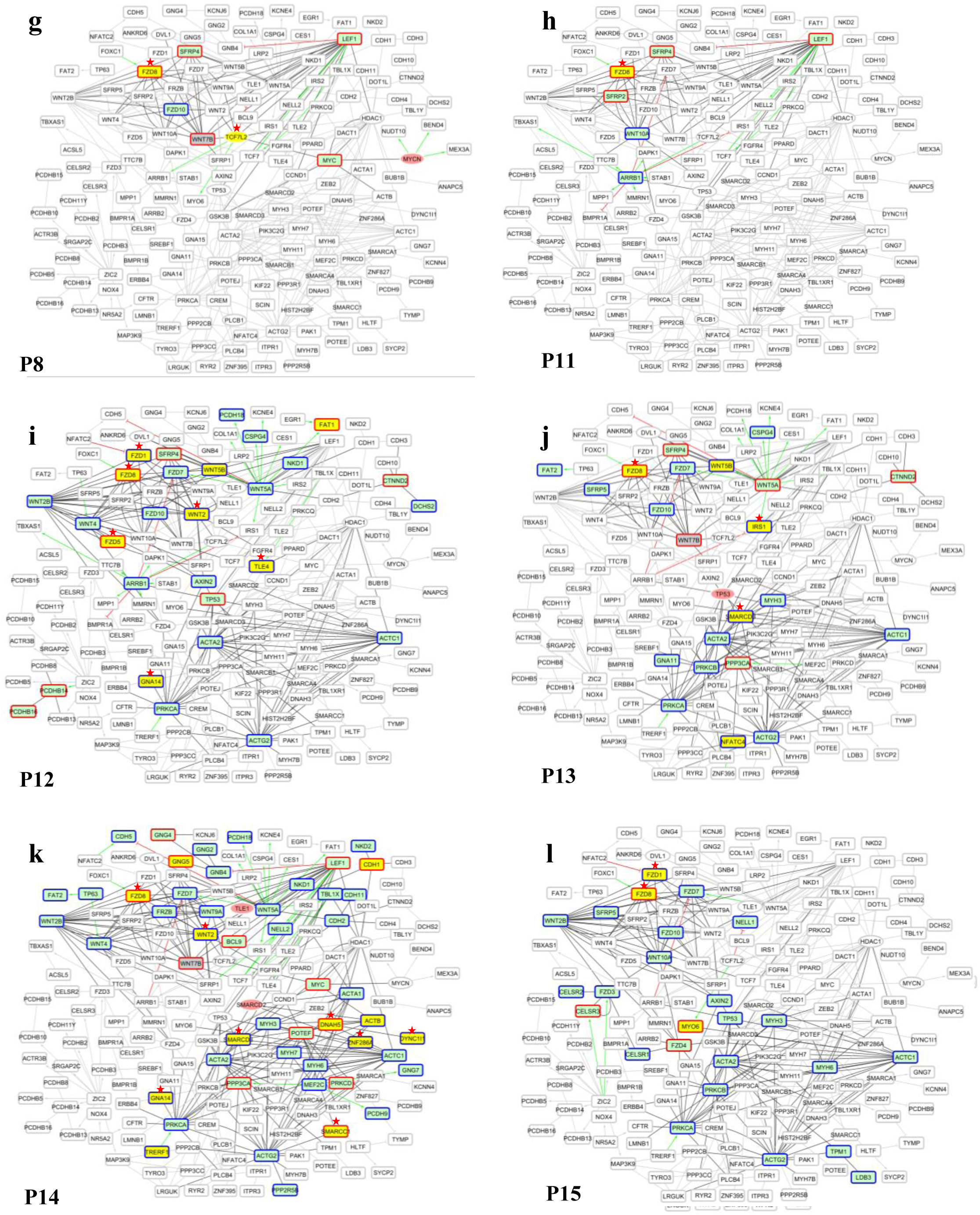
Patient-specific deregulations of Wnt pathway components for the indicated patients (P). **a,** Merged network of all 15 patients used a background in (b) to (l). Frame and color codes are shown below P5. **b** to **l**, deregulated and/or mutated components of the Wnt pathway in each patient.

**Extended Data Fig. 4:**
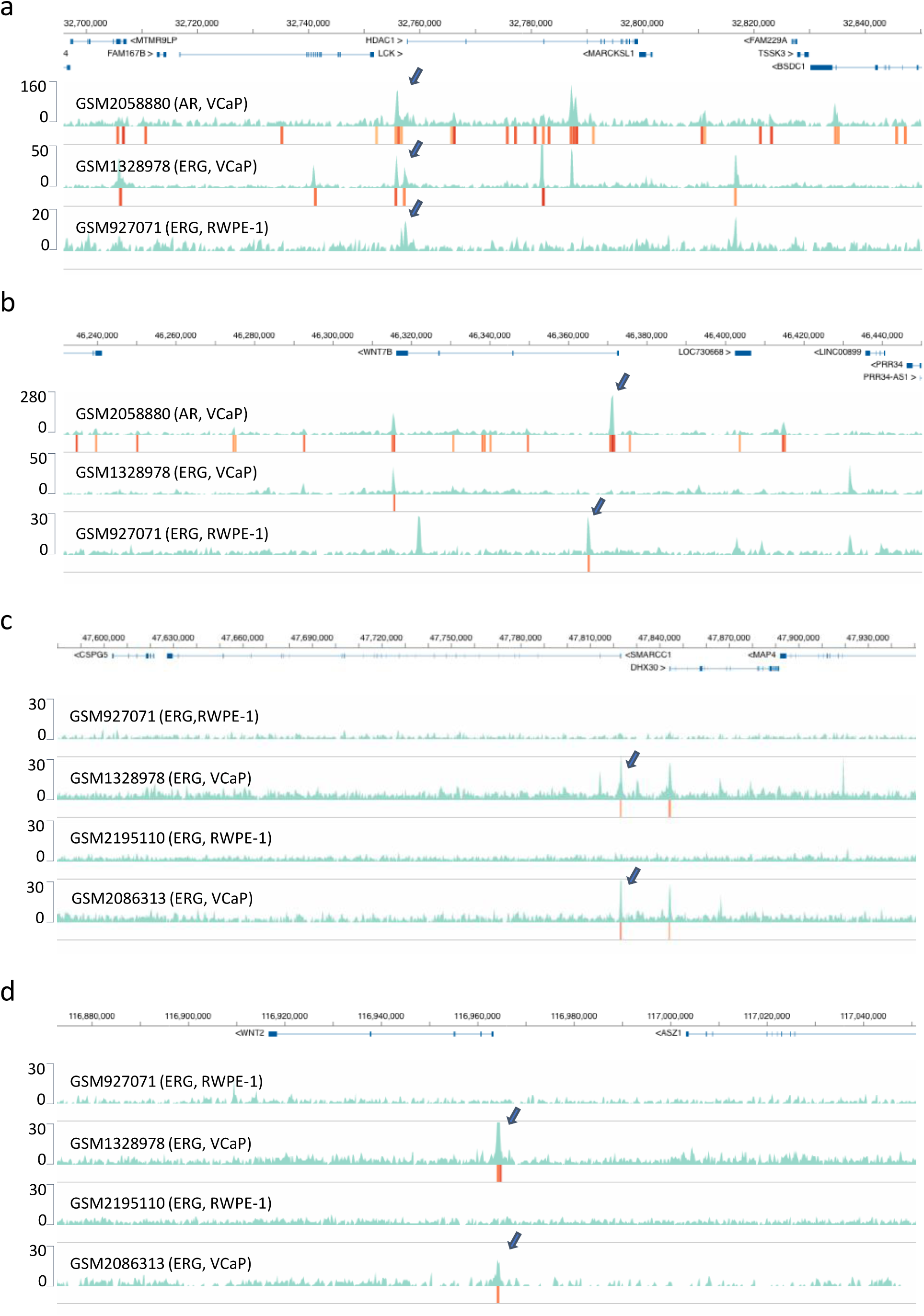
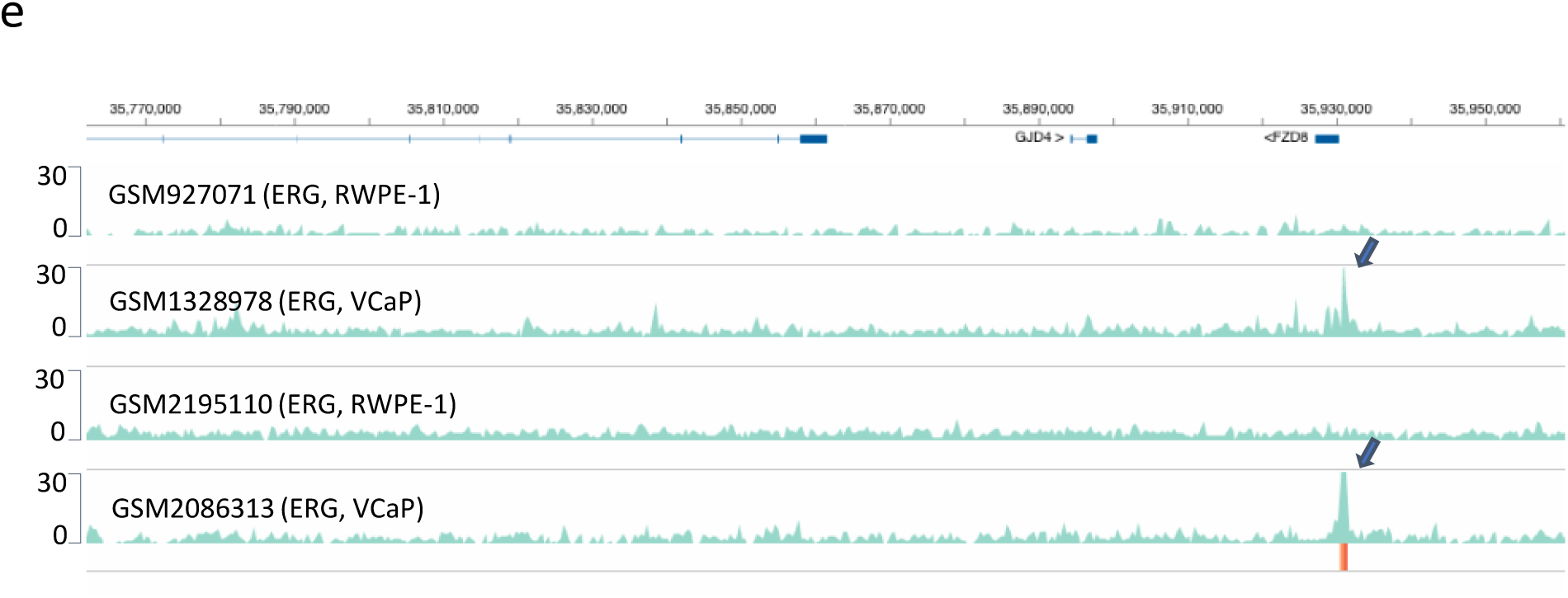
AR binding and acquisition of ERG binding sites in VCaP prostate cancer cells relative to RWPE-1 prostate epithelial cells. Screenshots of qcGenomics (http://www.ngs-qc.org/qcgenomics/) browser NAVi displaying genes that show AR and/or ERG binding in their promoter regions. **a,** *HDAC1*; **b,** *WNT7B;* **c,** *SMARCCl*; **d,** *WNT2*; **e,** *FDZ8*. ChlP-seq data sets in **a** and **b** are from GEO accession numbers (from top to bottom) GSM2058880 (AR, VCaP), GSM1328978 (ERG, VCaP) and GSM927071 (ERG, RWPE-1), as specified. Note that in (a) ERG binding at the *HDAC1* promoter is seen in VCaP and RWPE-1 cells, while in (b) for *WNT7B* a promoter-proximal ERG binding is seen in ‘normal’ RWPE-1 but not in VCaP cells; this ERG binding site is distant from the AR binding site. The ERG ChlP-seq data sets in **c, d** and **e** are from GEO accession numbers (from top to bottom) GSM927071 for RWPE-1, GSM1328978 for VCaP (both use anti-ERG antibody Epitomics 2805-1), GSM2195110 for RWPE-1 and GSM2086313 for VCaP. GSM2195110 was done by using Anti-ERG Clone 9FY Biocare # CM421 C, GSM2195110 used an anti-ERG antibody but did not provide the source. Note the consistency between corresponding experiments with different antibodies.

**Extended Data Fig. 5:**
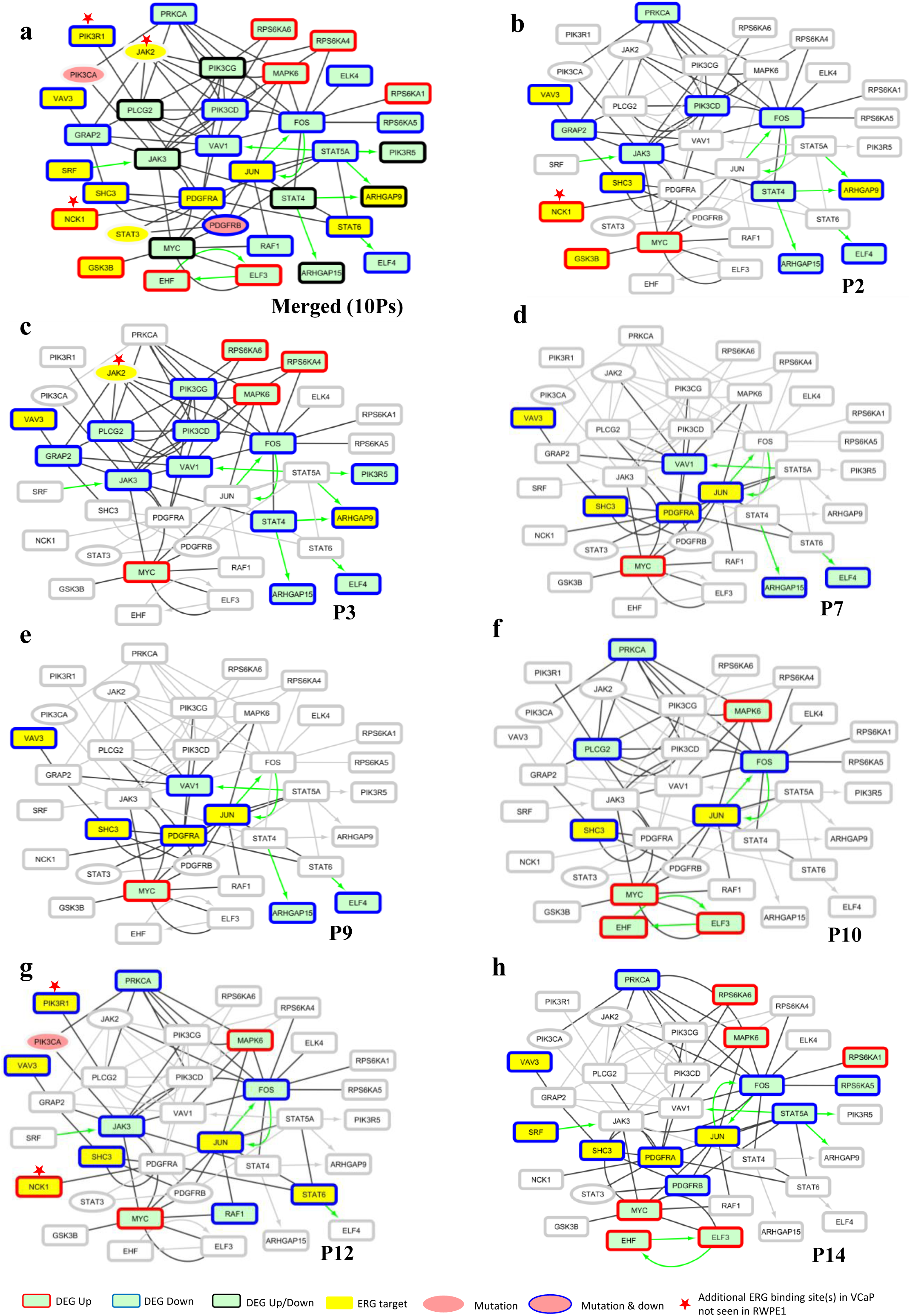
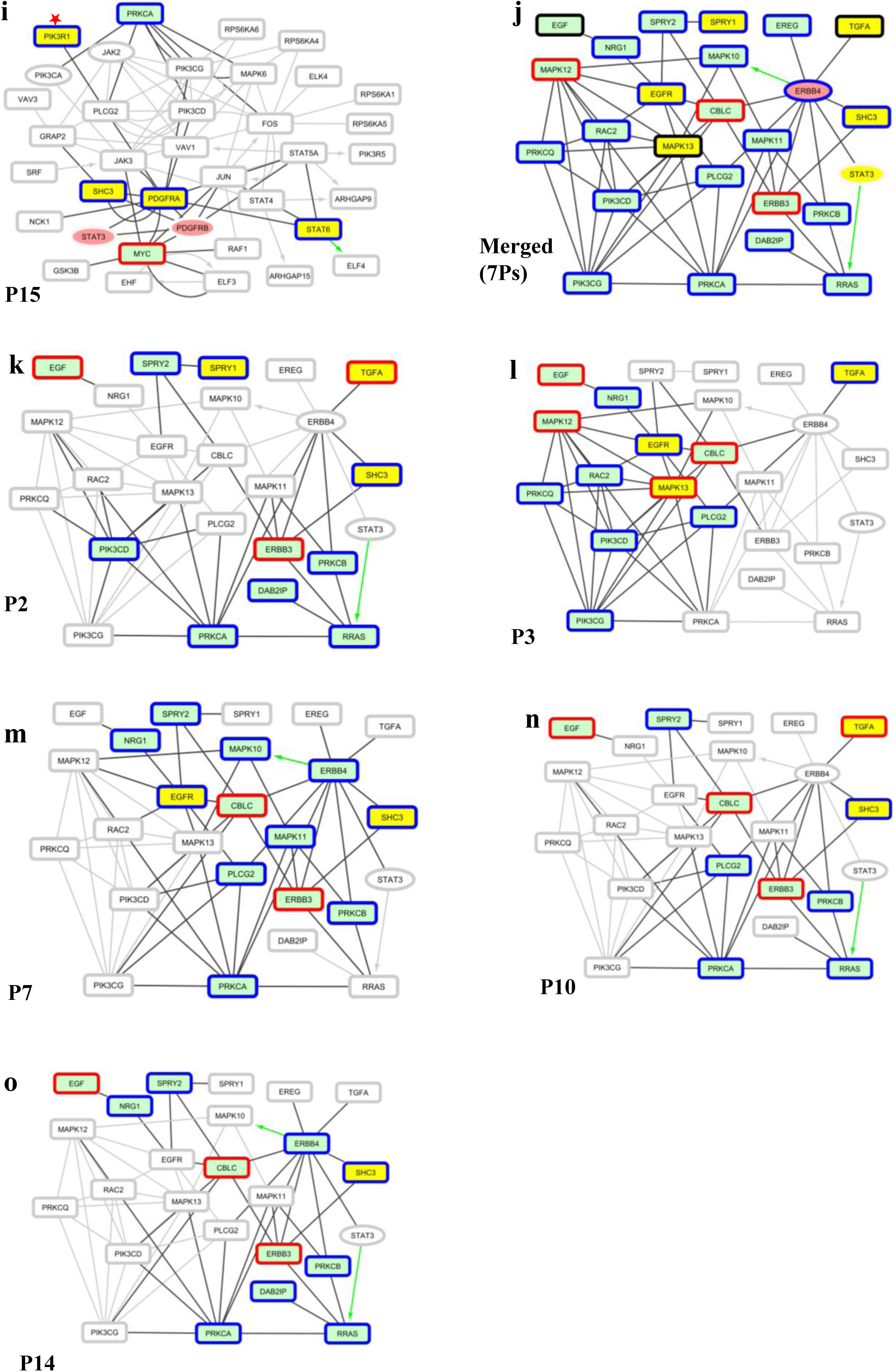
Patient-specific deregulations of the PDGF and EGFR pathway components for the indicated patients (P). **a,** Merged network of 10 patients used a background in (b) to (l) for PDGF pathway. **j,** Merged network of 7 patients used as a background in (k) to (o) for EGFR pathway. Frame and color codes are shown below P12. **b** to **i**, deregulated and/or mutated components of the PDGF pathway in each patient. **k** to **o** deregulated and/or mutated components of EGFR pathway in each patient.

**Extended Data Fig. 6:**
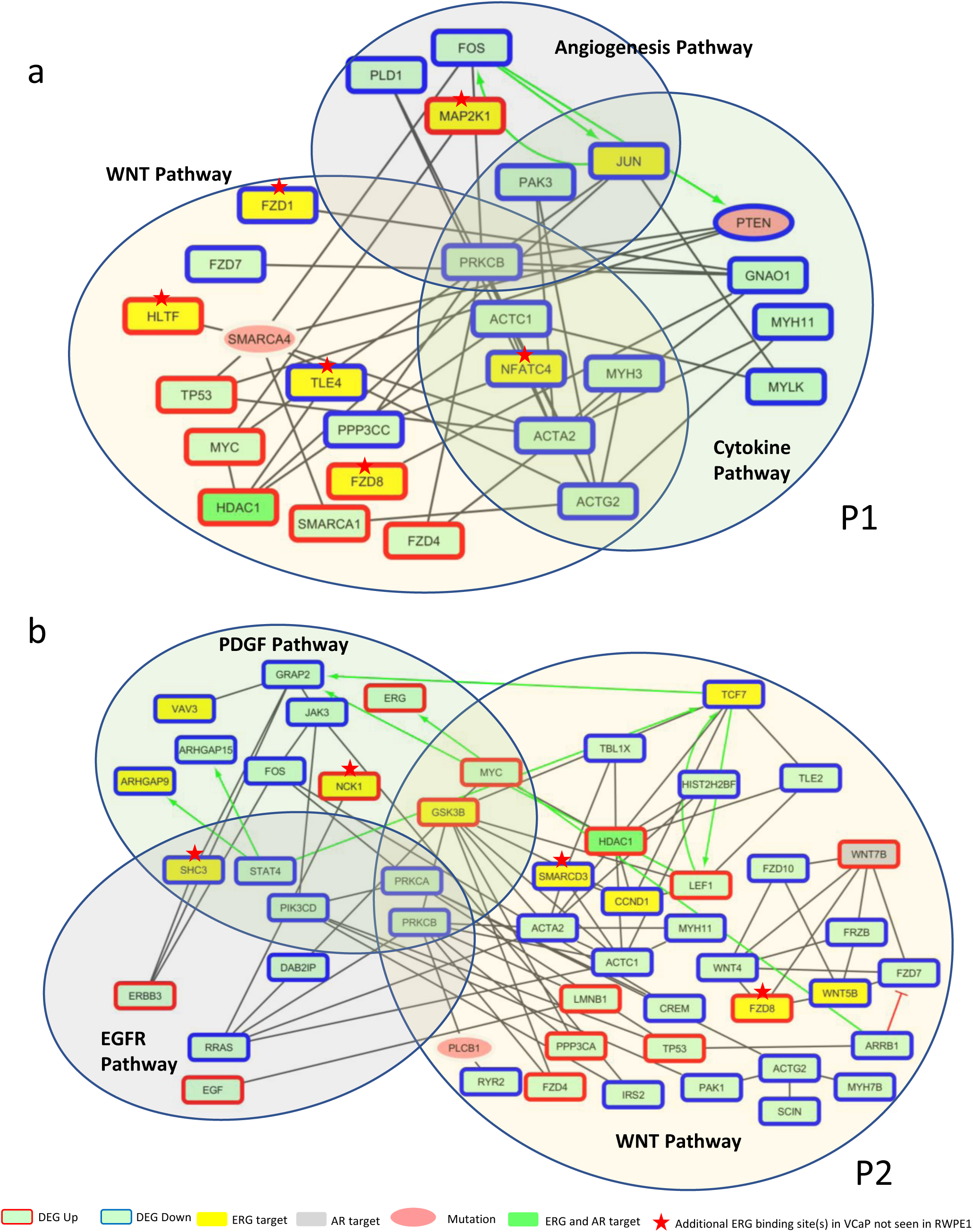
Patient-specific deregulations of the PDGF and EGFR pathway components for the indicated patients (P). **a,** Network display of deregulated and mutated factors of patient P1 in the WNT, Angiogenesis and Cytokine Pathways to reveal connectivities between the different pathways. **b,** Similar representation of the de-regulated and mutated factors in P2 for the WNT, PDGF and EGFR pathways.

## Methods

### Patient sample collection

Samples from non-treated patients were obtained after informed consent in accordance with the Declaration of Helsinki and stored at the « CRB Cancer des Hôpitaux de Toulouse (BB-0033-00014) » collection. According to the French law, CRB Cancer collection has been declared to the Ministry of Higher Education and Research (DC-2008-463) and obtained a transfer agreement (AC-2013-1955) after approbation by ethical committees. Clinical and biological annotations of the samples have been declared to the CNIL (Commission Nationale de l’Informatique et des Libertés). All samples were collected within 15 minutes after radical prostatectomy to shorten the delay between de-vascularization and freezing, and to ensure preservation of labile molecules. Immediately following prostatectomy, punch biopsies (“carrots”) of 8mm diameter were taken from tumor and adjacent normal tissue, snap frozen in liquid nitrogen and stored at −80 ° C. Carrots used for genomic and transcriptomic studies were cut into sequentially tissue sections and the tumor cellularity was monitored at regular intervals by histological staining to ensure homogeneity of tumor and normal sections stored in LoBind tubes at −80°C.

### Tissue microarrays

Tissues microarrays (TMA) were made from paraffin-embedded tissue cores of histopathologically-confirmed prostate cancer and patient-matched tumor-adjacent normal tissue. For each tumor, two representative tumor areas were selected and two cores of 2 mm in diameter were punched and included in paraffin recipient blocs. Two adjacent normal tissues of each selected prostatic sample were arrayed on TMAs and constituted the “normal” counterparts of each tumor sample. The TMAs were performed on the histopathology platform of the Biological Resource Center (CRB) Cancer of the Toulouse University Hospital, in a semi-automated way using the EZ-TMA™ Manual Tissue Microarray Kit, IHCWORLD. The slides were examined by HE coloration and immunohistochemical studies were performed on TMA tissues. Immunohistochemistry was done using an automated Dako Autostainer. The following antibodies were used: ERG (Epitomics, EPR3864, dilution 1/100, pH 6), EZH2 (Novocastra, NCL-L-EZH2, 1/50, pH 9); Androgen Receptor (Dako, M3562, 1/200, pH 9). Slides were digitalized using a Hamamatsu NanoZoomer slide scanner (Japan) at 20x magnification with a resolution of 0.46 microns per pixel. The results were interpreted under an optical microscope by two pathologists (CM and M-LQ), blinded to the clinical data.

### Whole exome sequencing (WES) and analysis pipeline

For WES, DNA was isolated from frozen tumor and matched normal tissue using QIAamp DNA micro kit (Cat No.56304 Qiagen) according to manufacturer’s instructions. DNA was processed by GATC Biotech for exome capture, library preparation and sequencing. Briefly, SureSelectXT Human all exon V6 kit (Agilent Technologies) was used to capture exons, libraries were prepared using TruSeq DNA library preparation kit (Illumina inc) according to manufacturer’s instruction and Paired-End 125-base sequencing was performed on Illumina HiSeq 2000. FastQ files provided by GATC Biotech were processed for variant discovery with Genome Analysis Toolkit (GATK, 3.7)^35^ using default parameters.

To assist in WES analysis we developed a WES Analysis Pipeline [written in Python3 with the Snakemake (3.13.3)^36^ management tool] and used the Genome Analysis Toolkit (GATK, 3.7)^35^ according to the authors’ instructions (download access in Supplementary Information). The following tools were used for each step in the pipeline:

*<u>Pre-processing of the samples</u>.* FastQ files were first aligned to hg19 (https://drive.google.com/open?id=1EuPivu7Quj5XiJlojpIxIREmyb4y3NCW, Genome Reference Consortium Human Build 37) using BWA-mem (0.7.17)^37^ using standard parameters. The output SAM files were then converted to BAM files using SAMtools (1.6)^38^. BAM files were processed using Picard tools (2.14) to sort by coordinates, remove duplicates and add read group tags (essential to differentiate between Normal and Tumor samples) to samples before indexing them with SAMtools. Then BAM files were recalibrated (BQSR) using GATK, as recommended for enhancing variant calling (https://gatkforums.broadinstitute.org/gatk/discussion/44/base-quality-score-recalibration-bqsr) by providing databases of known polymorphic sites: a set of curated INDEL entries (Mills_and_1000G_gold_standard.indels.hg19.sites.vcf, https://drive.google.com/open?id=1tXzEFrydWBzxWoP1iQS8QuYrXW2x75TU), a Single Nucleotide Polymorphism database dbSNP (dbsnp_138.hg19.vcf, https://drive.google.com/open?id=1d8OfwtB7zW8Tl27YxuFIF8eSkj3AgP_r), the COSMIC database of somatic cancer mutations (CosmicCodingMuts_GHCr37.p13.vcf, https://drive.google.com/open?id=1tXzEFrydWBzxWoP1iQS8QuYrXW2x75TU). *Creating the <u>Panel of Normals (PON)</u>.* The ‘Panel of Normals’ is created from the normal samples using GATK. This method is used as a filter to reject artifacts and germline variants that are present in at least two normal samples (-minN 2). It uses as input the hg19, the dbSNP, the COSMIC and the intervals of the genome to analyze only the exons of all genes captured (HumanAllExonV6r2, https://drive.google.com/open?id=1rxCn7x4SWAg9r8bYAmDzBV-8en-f8KLa). It generates a new file (https://drive.google.com/open?id=1MGuJYBTlzAUFKD0WfiEGwiU3MXUHgHpf) that will be used when calling the variants*. <u>Variant calling with MuTect2</u>*. After pre-processing and alignment, MuTect2 (GATK) was used to call somatic variants^39^. Inputs are the following files: hg19, PON, HumanAllExonV6r2, COSMIC and dbSNP. The normal and tumor samples were compared using the following parameters: pir_mad_threshold: 6; max_alt_alleles_in_normal_count: 5; pir_median_threshold: 35; standard_min_confidence_threshold_for_calling: 30. *<u>Annotation of VCF</u>.* The annotations of the VCF files were done using SnpEFF^40^ and SnpSift^40^ with hg19 as reference genome.

### Validation of mutations

Target regions were amplified by PCR. PCR products were purified using Qiagen gel extraction kit (28704) and sequenced by Eurofins Genomics using the BigDye Terminator Cycle Sequencing Kit and an ABI 3730xl automated sequencer (Applied Biosystems). The sequencing primers were the same as those used for PCR amplification. Variants were confirmed using SNAP gene viewer. For primer sequences, see Supplementary Information.

### RNA-sequencing

RNA was isolated from frozen tumor and matched normal tissue using Trizol reagent (Invitrogen). RNA was further cleaned up using RNEasy minElute RNA clean up kit (Qiagen). RNA was then sent to GATC Biotech (Konstanz, Germany) for strand specific, paired-end and Ribo-minus total RNA-seq. Briefly, ribosomal RNA depletion was done using Ribo Zero gold kit (Illumina Inc); libraries were prepared using TruSeq stranded total RNA library prep kit (Illumina Inc.). Paired-end 125 base or 150 base sequencing was performed using Illumina HiSeq 2000. FastQ files received from GATC Biotech were used for further analysis.

### RNA-seq analysis pipeline

The analysis pipeline consists of the following steps. *<u>Pre-processing, alignment and counting raw reads</u>.* FastQ files were assessed for quality using FastQC. FastQ files were aligned to reference genome (human genome hg19) using the Hisat2^41^ aligner. Aligned SAM files were converted to BAM files and sorted using SAMtools^38^. The R package SummarizedExperiment^42^ was used for counting raw reads per exon/gene.

*<u>Differential gene expression analysis.</u>* The patient specific differential gene expression analysis was done using DESeq2 (1.20.0)^43^ according to the general steps described with the parameters given below. The samples have been analyzed by giving the matched raw read counts normal/tumor duplicates as input.

− Removing sum of row counts: 0;

− CooksCutoff: False;

− Alpha: 0.01;

− Subset genes with Adjusted P-value ≤ 0.01;

− Subset genes with Log2FC ≤ -1 or Log2FC ≥ +1.

The corresponding list of Differentially Expressed Genes (DEGs) for each patient was used for further analysis.

### Pathway enrichment analysis

To interpret the gene expression data, the DEG list was loaded into GeneCodis^30^ and the Panther pathway analysis function was used to retrieve enriched pathways. Hypergeometric correction of p-values was applied and pathways displaying a corrected p-value < 0.01 were considered enriched. We clustered all pathways of all samples using Plotly (4.8.0) (https://plot.ly/) in R. For the datasets obtained from TCGA (https://portal.gdc.cancer.gov/, 54 PrCa patient data along with matched normal), HT-seq counts were downloaded for each patient corresponding to tumor and matched normal. DEseq2 was used to identify the DEGs for each patient. Pathway enrichment analysis was performed in as described before.

### Patient-specific network generation and visualization

To generate the gene networks for individual patients we extracted the list of mutated genes from WES and differentially expressed genes (DEGs) from RNA-seq of tumor *vs* normal samples for each patient. These lists of genes were queried against two known databases of network interactions, STRING^44^, a Protein-Protein Interaction (PPI) database, and CellNet^7^, a gene regulatory network (GRN) database. For STRING, we merged the list of genes (DEGs and Mutation, keeping the information whether the gene is a DEG or a mutated gene as attributes), removed any duplicated genes and queried them using an in-house script (Supplementary Information). As for the parameters, we only chose edge interactions that have been experimentally validated (exp_score ≠ 0). For CellNet, we queried only the differentially expressed genes on the target genes and retrieved along the cognate transcription factors. We chose interactions who had only a z-score ≥ 5. After obtaining networks from both databases, we proceed to add the information from WES and RNA-seq whether the genes were mutated, differential expressed or both, in addition to the information obtained from the databases.

### Network visualization and merging using Cytoscape

Individual networks, created by using Cellnet and String for each patient, were visualized using Cytoscape^45^. Finally, CellNet and STRING networks for each patient were merged using the Cytoscape merge function to obtain master networks for each patient. Sub-networks were then extracted for further visualization and analysis.

### Identification of putative AR and ERG target genes

A two-step approach was used. First, we collected sequenced read files (bed format) associated to public ChIP-seq assays targeting ERG in TMPRSS2-ERG positive human VCaP prostate cancer (GSM1328978, GSM1328979) and RWPE-1 normal prostate epithelium cells (GSM2195103, GSM2195106). BED Replicate files per cell-type were merged together prior performing peak calling (MACS 1.4; no model, shiftsize=150nts, p-value threshold: 1×10^−5^), followed by their genomic annotation to the closest transcription start sites (<u>annoPeakR</u>). This analysis allowed to pair the characterized DEGs and mutated genes within the patient-derived networks with genes presenting proximal AR binding sites (<10 kb distance) on VCaP ChIP-seq profiles. This primary analysis has been validated in a second step by comparative visual inspection of ChIP-seq profiles. For this we used the qcGenomics platform (http://ngs-qc.org/qcgenomics/), in which the dedicated genome browser NAVi allows to visualize any publicly available ChIP-seq profile. Specifically, we used NAVi to extract all AR and ERG ChIP-seq profiles for TMPRSS2-ERG positive human VCaP prostate cancer and RWPE1 normal prostate epithelium cells. The pre-computed datasets were displayed simultaneously in the NAVi browser for comparative visualization. Only tracks with an apparent high signal-to-noise ratio were retained (VCaP-ERG: GSM2058880, GSM1328978, GSM1378979, GSM1328980, GSM1328981; VCaP-AR: GSM1410768, RWEP1-ERG: GSM927071, GSM2195110, GSM2195103; VCaP-GROseq: GSM2235682). Promoter-proximal ERG binding was scored positive in this visual ‘validation’ (attributing a yellow color to the respective nodes) only when there was a clearly visible peak above the background at a scale of 30 to 300 (read count intensity; depending on the signal and noise intensities of each profile), provided that there was no other known TSS closer (see Extended Data Fig. 4 for examples of gain of ERG binding).

### Oligonucleotide sequences

Primers used for PCR are specified in Supplementary Table 5 of the Supplementary Information.

### Data availability

The RNA-seq data sets generated in the context of this study from 15 patient-matched tumor and normal prostate tissue are available in the Gene Expression Omnibus (GEO) repository under the accession number GSE133626. The corresponding Exome-seq data sets from the prostates of the same 15 patients are available from the SRA database under the accession number [Data sets have been submitted, accession numbers will be provided when attributed]. Data sets for the pathway analysis shown in Extended Data Fig. 2 were downloaded from The Cancer Genome Atlas (TGCA) as described in the methods section. The sequencing statistics of RNA-seq and Exome-seq experiments are specified in Supplementary Table 4 of the Supplementary Information.

## References

1. Bell, K.J., Del Mar, C., Wright, G., Dickinson, J. & Glasziou, P. Prevalence of incidental prostate cancer: A systematic review of autopsy studies. Int J Cancer 137, 1749–57 (2015).

2. Robinson, D. et al. Integrative clinical genomics of advanced prostate cancer. Cell 161, 1215–1228 (2015).

3. Farashi, S., Kryza, T., Clements, J. & Batra, J. Post-GWAS in prostate cancer: from genetic association to biological contribution. Nat Rev Cancer 19, 46–59 (2019).

4. Abida, W. et al. Genomic correlates of clinical outcome in advanced prostate cancer. Proc Natl Acad Sci U S A 116, 11428–11436 (2019).

5. Drake, J.M. et al. Phosphoproteome Integration Reveals Patient-Specific Networks in Prostate Cancer. Cell 166, 1041–1054 (2016).

6. Szklarczyk, D., et al. STRING v11: protein-protein association networks with increased coverage, supporting functional discovery in genome-wide experimental datasets. Nucleic Acids Res (2018).

7. Cahan, P. et al. CellNet: network biology applied to stem cell engineering. Cell 158, 903–15 (2014).

8. Network, T.C.G.A.R. The Molecular Taxonomy of Primary Prostate Cancer. Vol. 163 1011–1025 (Cell, 2015).

9. Baca, S.C. et al. Punctuated evolution of prostate cancer genomes. Cell 153, 666–77 (2013).

10. Tomlins, S.A. et al. Recurrent fusion of TMPRSS2 and ETS transcription factor genes in prostate cancer. Science 310, 644–8 (2005).

11. Chen, C.D. et al. Molecular determinants of resistance to antiandrogen therapy. Nat Med 10, 33–9 (2004).

12. Haffner, M.C. et al. Tracking the clonal origin of lethal prostate cancer. J Clin Invest 123, 4918–22 (2013).

13. Yi, S. et al. Functional variomics and network perturbation: connecting genotype to phenotype in cancer. Nat Rev Genet 18, 395–410 (2017).

14. Kypta, R.M. & Waxman, J. Wnt/β-catenin signalling in prostate cancer. Nat Rev Urol 9, 418–28 (2012).

15. Jalota, A. et al. Tumor suppressor SMAR1 activates and stabilizes p53 through its arginine-serine-rich motif. J Biol Chem 280, 16019–29 (2005).

16. Ishii, H. et al. FEZ1/LZTS1 gene at 8p22 suppresses cancer cell growth and regulates mitosis. Proc Natl Acad Sci U S A 98, 10374–9 (2001).

17. Vecchione, A. et al. Fez1/Lzts1 absence impairs Cdk1/Cdc25C interaction during mitosis and predisposes mice to cancer development. Cancer Cell 11, 275–89 (2007).

18. Shen, M. et al. Tinagl1 Suppresses Triple-Negative Breast Cancer Progression and Metastasis by Simultaneously Inhibiting Integrin/FAK and EGFR Signaling. Cancer Cell 35, 64–80.e7 (2019).

19. Armenia, J. et al. The long tail of oncogenic drivers in prostate cancer. Nat Genet 50, 645–651 (2018).

20. Jin, Y. et al. Molecular circuit involving KLK4 integrates androgen and mTOR signaling in prostate cancer. Proc Natl Acad Sci U S A 110, E2572–81 (2013).

21. Ramsay, A.J. et al. Kallikrein-related peptidase 4 (KLK4) initiates intracellular signaling via protease-activated receptors (PARs). KLK4 and PAR-2 are co-expressed during prostate cancer progression. J Biol Chem 283, 12293–304 (2008).

22. Adriaens, C. et al. p53 induces formation of NEAT1 lncRNA-containing paraspeckles that modulate replication stress response and chemosensitivity. Nat Med 22, 861–8 (2016).

23. Mello, S.S. et al. is a p53-inducible lincRNA essential for transformation suppression. Genes Dev 31, 1095–1108 (2017).

24. Zhang, Y. et al. Analysis of the androgen receptor-regulated lncRNA landscape identifies a role for ARLNC1 in prostate cancer progression. Nat Genet 50, 814–824 (2018).

25. Malek, R. et al. TWIST1-WDR5-. Cancer Res 77, 3181–3193 (2017).

26. Malik, R. et al. Targeting the MLL complex in castration-resistant prostate cancer. Nat Med 21, 344–52 (2015).

27. Rupaimoole, R. & Slack, F.J. MicroRNA therapeutics: towards a new era for the management of cancer and other diseases. Nat Rev Drug Discov 16, 203–222 (2017).

28. Garofalo, M. et al. miR-221&222 Regulate TRAIL Resistance and Enhance Tumorigenicity through PTEN and TIMP3 Downregulation. Cancer Cell 16, 498–509 (2009).

29. Mi, H., Muruganujan, A., Ebert, D., Huang, X. & Thomas, P.D. PANTHER version 14: more genomes, a new PANTHER GO-slim and improvements in enrichment analysis tools. Nucleic Acids Res 47, D419–D426 (2019).

30. Tabas-Madrid, D., Nogales-Cadenas, R. & Pascual-Montano, A. GeneCodis3: a non-redundant and modular enrichment analysis tool for functional genomics. Nucleic Acids Res 40, W478–83 (2012).

31. Lunardi, A. et al. Suppression of CHK1 by ETS Family Members Promotes DNA Damage Response Bypass and Tumorigenesis. Cancer Discov 5, 550–63 (2015).

32. Kahn, M. Can we safely target the WNT pathway? Nat Rev Drug Discov 13, 513–32 (2014).

33. Chen, S. et al. Widespread and Functional RNA Circularization in Localized Prostate Cancer. Cell 176, 831–843 e22 (2019).

34. Vo, J.N. et al. The Landscape of Circular RNA in Cancer. Cell 176, 869–881 e13 (2019).

## References

35. McKenna, A. et al. The Genome Analysis Toolkit: a MapReduce framework for analyzing next-generation DNA sequencing data. Genome Res 20, 1297–303 (2010).

36. Koster, J. & Rahmann, S. Snakemake--a scalable bioinformatics workflow engine. Bioinformatics 28, 2520–2 (2012).

37. Li, H. & Durbin, R. Fast and accurate short read alignment with Burrows-Wheeler transform. Bioinformatics 25, 1754–60 (2009).

38. Li, H. et al. The Sequence Alignment/Map format and SAMtools. Bioinformatics 25, 2078–9 (2009).

39. Cibulskis, K. et al. Sensitive detection of somatic point mutations in impure and heterogeneous cancer samples. Nat Biotechnol 31, 213–9 (2013).

40. Cingolani, P. et al. A program for annotating and predicting the effects of single nucleotide polymorphisms, SnpEff: SNPs in the genome of Drosophila melanogaster strain w1118; iso-2; iso-3. Fly (Austin) 6, 80–92 (2012).

41. Kim, D., Langmead, B. & Salzberg, S.L. HISAT: a fast spliced aligner with low memory requirements. Nat Methods 12, 357–60 (2015).

42. Lawrence, M. et al. Software for computing and annotating genomic ranges. PLoS Comput Biol 9, e1003118 (2013).

43. Love, M.I., Huber, W. & Anders, S. Moderated estimation of fold change and dispersion for RNA-seq data with DESeq2. Genome Biol 15, 550 (2014).

44. Szklarczyk, D. et al. The STRING database in 2017: quality-controlled protein-protein association networks, made broadly accessible. Nucleic Acids Res 45, D362–D368 (2017).

45. Shannon, P. et al. Cytoscape: a software environment for integrated models of biomolecular interaction networks. Genome Res 13, 2498–504 (2003).

