## Supplementary information for "Patient-matched analysis identifies deregulated networks in prostate cancer to guide personalized therapeutic intervention"

**Patient-matched analysis to identify perturbed gene-regulatory networks and therapeutic targets in advanced prostate cancer**

**Akinchan Kumar^1,2,3,4,5^, Alaa Badredine^1,2,3,4,5^, Karim Azzag^6^, Yasenya Kasikçi^1,2,3,4,5^,**

**Marie Laure Quintyn Ranty^7^, Falek Zaidi^7^, Nathalie Serret^7^, Catherine Mazerolles^7^, Bernard Malavaud^7^, Marco Antonio Mendoza-Parra^1,2,3,4,5^, Laurence Vandel^6^ and Hinrich Gronemeyer^1,2,3,4,5^**

^1^ Institut de Génétique et de Biologie Moléculaire et Cellulaire (IGBMC), Department of Functional Genomics and Cancer, Illkirch, France ; ^2^ Centre National de la Recherche Scientifique, UMR7104, Illkirch, France ; ^3^ Institut National de la Santé et de la Recherche Médicale, U1258, Illkirch, France ; ^4^ Université de Strasbourg, Illkirch, France ; ^5^ Equipe Labellisée Ligue Contre le Cancer ; ^6^ Centre de Biologie du Développement (CBD), Centre de Biologie Intégrative (CBI), Université de Toulouse, CNRS, UPS, France ; ^7^ Institut Universitaire du Cancer Toulouse-Oncopole (IUCT-O), Toulouse, France

### Supplementary Table 1: Number of mutations and differentially expressed genes in each patient

| Patient N° (this study) | N° of passed mutations | N° of moderate mutations | N° of high mutations | N° of DEGs  (FC > 2; adj p -value < 0.01) |
| --- | --- | --- | --- | --- |
| 1 | 86 | 34 | 9 | 1982 |
| 2 | 105 | 39 | 5 | 3170 |
| 3 | 71 | 29 | 7 | 2808 |
| 4 | 68 | 21 | 5 | 2652 |
| 5 | 49 | 20 | 6 | 4029 |
| 6 | 53 | 18 | 6 | 1034 |
| 7 | 87 | 31 | 8 | 2889 |
| 8 | 77 | 31 | 6 | 1439 |
| 9 | 83 | 27 | 3 | 3528 |
| 10 | 55 | 21 | 1 | 2159 |
| 11 | 83 | 31 | 11 | 1215 |
| 12 | 114 | 43 | 16 | 3438 |
| 13 | 89 | 27 | 6 | 1797 |
| 14 | 63 | 21 | 2 | 3563 |
| 15 | 68 | 24 | 2 | 2453 |

Patient samples that complied with the histological selection criteria for homogeneity and tumor cellularity were renumbered from 1 to 15. **Mutations.** Mutations were classified as defined by Ensembl (https://www.ensembl.org/info/genome/variation/prediction/predicted_data.html). **Differentially Expressed Genes (DEGs).** Duplicate RNA-seq of tumor and patient-matched normal tissue was used to define differentially expressed genes (DEGs) by counting the reads with Genomic Alignments (1.17.3) and then using DESeq2 (1.17.3) with R (3.5.0). The results have been obtained following the general guidelines given in the DESeq2 tool. We used a Log2FC of -1/+1 (+2/-2) and a cutoff for the adjusted p-value (Q-value) < 0.01. All main figures have been generated using the high, moderate and modifier mutations specified in Supplementary File 1.

### Supplementary Table 2: Novel mutations previously not reported for prostate cancer patients

| **Patient 1** | | Total number of somatic mutations: 86 | |
| --- | --- | --- | --- |
| **Gene** | **Type of Variant** | **New Mutation** | **Function** |
| *OR10H1* | Missense | T57M | olfactory receptor |
| *CDHR5* | Splicing mut | Splice donor exon 6 | member of the cadherin superfamily; enriched in colorectal cancer, liver cancer, pancreatic cancer, renal cancer, stomach cancer |
| *HSD17B6* | Missense | R52Q | 17 Beta-hydroxysteroid dehydrogenase type 6; involved in intra-tumoral synthesis of DHT |
| *TRMT10A* | Missense | R275Q | tRNA (Guanine-1)-methyltransferase |
| *BANP* | Missense | R246H | forms complex with p53 & negatively regulates p53 transcription, tumor suppressor and cell cycle regulator |
| *DGAT2* | Missense | L91R | Diacylglycerol O-Acyltransferase 2 |
| *CASD1* | Missense | I755S | O-acetyltransferase |
| *TTC28* | Frameshift | L1164LLTC….. | During mitosis, may be involved in the condensation of spindle midzone microtubules, leading to the formation of midbody |
| *ITGB7* | Frameshift & splice region | PCERHR644PGLCR….. | Integrin Subunit Beta 7 (ECM signaling) |
| *FEZ1* | Stop gain | E65* | leucine-zipper protein, its expression is altered in multiple human tumors; candidate tumor suppressor |
| *TINAGL1* | Missense | A424V | Inhibits TNBC progression and metastasis; prognostic marker in [renal cancer](https://www.proteinatlas.org/ENSG00000142910-TINAGL1/pathology/tissue/renal+cancer) (favourable) and [thyroid cancer](https://www.proteinatlas.org/ENSG00000142910-TINAGL1/pathology/tissue/thyroid+cancer) (favourable) |
| *ZFR2* | Missense | Q632H | Zinc-finger RNA binding protein |
| *SLC35B2* | Missense | S218F | Predicted membrane protein Transporter; Prognostic marker in liver (unfavorable), prostate (favorable) & [breast cancer](https://www.proteinatlas.org/ENSG00000157593-SLC35B2/pathology/tissue/breast+cancer) (unfavorable) |
| *PPP2R5D* | Missense | L183Q, | protein phosphatase 2 regulatory subunit B Delta isoform |
| *WDFY1* | Missense | L149P | phosphatidylinositol 3-phosphate binding protein (signaling) |

| **Patient 2** | | Total number of somatic mutations: 105 | |
| --- | --- | --- | --- |
| **Gene** | **Type of Variant** | **New Mutation** | **Function** |
| *NBR1* | Missense | V96I | Autophagy cargo receptor |
| *RHPN2* | Missense | V73M | Rhophilin-Like Rho-GTPase Binding Protein |
| *SH3PXD2A* | Missense | T847P | Adapter protein adhesion |
| *MUC2* | Missense | T2305A | Member of mucin family |
| *CHN2* | Missense | R47W | Role in cell proliferation and migration |
| *CCDC169* | Missense | R197T | Coiled-Coil Domain-Containing Protein 169 |
| *SPTLC2* | Frameshift Ins | P467SSS…, | Serine palmitoyltransferase |
| *TRIM49C* | Missense | K363E | Uncharacterized |
| *BRF1* | Missense | K282Q | RNA Polymerase III Transcription Initiation Factor Subunit |
| *HIST1H3J* | Missense | H114Y | AR signaling |
| *ZBTB42* | Missense | F82C | Transcriptional repressor |
| *LRRCC1* | Missense | E525K | Leucine Rich Repeat & Coiled-Coil Centrosomal Protein 1 |
| *ZNF44* | Missense | E348A | Uncharacterized |
| *PPP1R26* | Missense | C186F | Phosphatase |

| **Patient 4** | | Total number of somatic mutations: 68 | |
| --- | --- | --- | --- |
| **Gene** | **Type of Variant** | **New Mutation** | **Function** |
| *DERL3* | Missense | V106I | Degrades misfolded glycoproteins in the ER |
| *PDGFB* | Missense | R217W | catalytic receptors with intracellular tyrosine kinase activity. Regulate embryonic development, angiogenesis, cell proliferation and differentiation. |
| *TEKT4* | Missense & Stop gain | L366V | Contributes to sperm motility |
| *TET3* | Missense | L1035M | Methylcytosine Dioxygenase |
| *PRH2* | Missense | I42L | Inhibitor of calcium phosphates |
| *TAS1R3* | Missense | G314D | G-protein coupled receptor |
| *KRTAP3-2* | Frameshift del | A16APP… | Keratin associated protein |
| *DRG2* | Missense | A135T | GTP binding protein |
| *NPFF* | Missense | R21H | Morphine modulating peptide |

| **Patient 3** | | Total number of somatic mutations: 71 | |
| --- | --- | --- | --- |
| **Gene** | **Type of Variant** | **New Mutation** | **Function** |
| *GSG1L2* | Missense | V72L | Germ Cell-Specific Gene 1-Like Protein 2-Like, Uncharacterized |
| *ASB15* | Missense | V358I | Participate in the ubiquitin-proteasome system for the degradation of proteins in the cell cycle and signal transduction pathways |
| *SH3PXD2A* | Missense | T847P | Role in cell migration |
| *TOPAZ1* | Missense | R85H | Required for progression to the post-meiotic stages of spermatocyte development |
| *ZNF280A* | Missense | R457Q | Uncharacterized |
| *OTOG* | Missense | R2294C | Component of the acellular membranes of the inner ear |
| *LTK* | Missense | R221H | Tyrosine kinase |
| *LRRTM4* | Missense | Q87K | Development and maintenance of the vertebrate nervous system |
| *JAK2* | Missense | P121L | Tyrosine kinase |
| *RGPD2* | Missense | L12V | RNA transport |
| *KRT5* | in-frame del | GGGLGGGLA517G | Member of keratin gene family |
| *TEAD4* | Missense | I242L | Hippo signaling, EMT |
| *TTC34* | Missense | G538S | Uncharacterized |
| *CCDC71L* | Missense | G28V | Uncharacterized |
| *DUOXA1* | Missense | F130L | Involved in hydrogen peroxide production necessary for thyroid hormonogenesis |
| *VWCE* | Missense | A58V | Regulatory element in the beta-catenin signaling pathway |

| **Patient 5** | | Total number of somatic mutations: 49 | |
| --- | --- | --- | --- |
| **Gene** | **Type of Variant** | **New Mutation** | **Function** |
| *SCCPDH* | Missense & splice region | Y102C | Oxidoreductase |
| *WRN* | Missense | V1155L | Helicase |
| *ZBED5* | Missense & Stop gain | Q329* | Not well characterized |
| *HDAC1* | Missense | R77S | Histone deacetylase |
| *TPSB2* | Missense | R22P | Serine protease |
| *ZNF777* | Missense | P158A | Not well characterized |
| *IL15* | Start lost | M1L | Cytokine activity |
| *FEZ2* | Missense | K278E | Involved in axonal outgrowth |
| *FKBP9* | Missense | H620Q | Calcium ion binding |
| *IFITM2* | frameshift del | R86RRW | Cytokine signaling |
| *PIM3* | in-frame del | AT226 | Ser/thr kinase |

| **Patient 6** | | Total number of somatic mutations: 53 | |
| --- | --- | --- | --- |
| **Gene** | **Type of Variant** | **New Mutation** | **Function** |
| *ZP1* | Missense | Y271S | Structural component of zona pellucida |
| *FASTKD3* | Missense | R216H | Kinase |
| *MAST3* | Missense | M786V | Kinase |
| *TGFB3* | Missense & splice region | E119K | TGF beta/Smad signaling network |
| *DLL1* | Stop gain | C44* | Notch ligand |
| *ZBED8* | Missense | A523S | Uncharacterized |

| **Patient 7** | | Total number of somatic mutations: 87 | |
| --- | --- | --- | --- |
| **Gene** | **Type of Variant** | **New Mutation** | **Function** |
| *COLCA2* | Missense & splice region | T34N | Colorectal Cancer Associated 2 |
| *FASTKD3* | Missense | R216H | Kinase |
| *ATG101* | Missense | T133K | Autophagy |
| *HTATIP2* | Missense | R49S | Oxidoreductase required for tumor suppression |
| *ALDH2* | Missense | R338Q | Oxidoreductase |
| *TTC29* | Frameshift | N499K* | Uncharacterized |
| *ARHGAP45* | Missense | L371Q | GTPase activator |
| *EMC9* | Missense | F30Y | ER membrane protein |
| *CCDC168* | Missense | D5448G | Uncharacterized |
| *HLA-DRB1* | Missense | D31G | Antigen presentation |
| *SLC5A9* | Missense | A386V | Glucose transport |
| *LRRC18* | Missense | A217T | Sperm maturation |
| *SHROOM2* | Missense | A1406V | Endothelial cell morphology |

| **Patient 8** | | Total number of somatic mutations: 77 | |
| --- | --- | --- | --- |
| **Gene** | **Type of Variant** | **New Mutation** | **Function** |
| *ZNF354C* | Missense | V176I | Transcriptional repressor |
| *ZMYM6* | Missense | S300C | Regulates cell morphology |
| *RABGGTB* | Frameshift & stop gain | R65RMN…. | RAB GTPase Binding |
| *ALDH2* | Missense | R338Q | Oxidoreductase |
| *SCRN1* | Missense | R50W | Peptidase |
| *RASSF8* | Missense | R208H | Tumor suppressor |
| *ITPRIPL2* | Missense | P171A | Intracellular calcium signaling |
| *ZBTB14* | Missense | H355Q | Not well characterized |
| *CYB5A* | Missense | F63V | Cytochrome C oxidase |
| *EFNA1* | Missense | E55A | Receptor tyrosine kinase |
| *SLC11A1* | Missense | A505S | Fe and Mn transporter |
| *OR5M9* | Missense | A298P | Signaling by GPCR |

| **Patient 9** | | Total number of somatic mutations: 83 | |
| --- | --- | --- | --- |
| **Gene** | **Type of Variant** | **New Mutation** | **Function** |
| *METTL4* | Missense | Y79F | Probable methyltransferase |
| *ROGDI* | Missense | V244M | Uncharacterized |
| *ZIC2* | Missense & in-frame del | S422C | Transcription repressor/activator |
| *MAPK7* | Missense | R400H | Role in proliferation |
| *AMIGO3* | Missense | R293W | May regulate cell-cell interaction |
| *GABPB2* | Missense | R117Q | May function as TF for purine rich repeats |
| *OLFM3* | Missense | N251S | Uncharacterized |
| *OR5AP2* | Missense | I220T | Olfactory receptor |
| *DBR1* | in-frame del | DD541D | RNA lariat debranching enzyme that hydrolyzes 2'-5' prime branched phosphodiester bonds |
| *GAK* | Missense & splice region | D1016Y | Cyclin G associated kinase |
| *GOPC* | Missense | A203V | Protein trafficking |

| **Patient 10** | | Total number of somatic mutations: 55 | |
| --- | --- | --- | --- |
| **Gene** | **Type of Variant** | **New Mutation** | **Function** |
| *ACSM4* | Missense | R325W | Fatty acid beta-oxidation |
| *RAB29* | Missense | P131A | Rab GTPase |
| *PPP1R10* | Missense & in-frame del | H896Q | Phosphatase |
| *TBL2* | Missense | H400Y | Beta transducin |
| *EHD3* | Missense | F18L | Membrane re-organization |
| *PMEL* | Missense | T419K | Melanosome biogenesis |
| *DNAH12* | Missense | A1501T | Dynein |

| **Patient 11** | | Total number of somatic mutations: 83 | |
| --- | --- | --- | --- |
| **Gene** | **Type of Variant** | **New Mutation** | **Function** |
| *TMEM132C* | Missense | V51G | Protein phosphatase |
| *SRM* | Missense | V112M | Spermidine biosynthesis |
| *WHRN* | Missense | S175T | Actin cytoskeletal assembly |
| *ADAMTS4* | Missense | R447H | Metalloproteinase |
| *SIGLEC12* | Missense | P198L | Protein carbohydrate interaction |
| *ANXA1* | Frameshift del | aa315* | Tumor suppressor. Inhibits phospholipase A2; anti-inflammatory |
| *STEAP1* | Frameshift del | aa244* | Cell surface antigen expressed at cell-cell junctions |
| *OR4E2* | Frameshift del | aa147* | Olfactory receptor |
| *ZFP30* | Missense | E159G | Not well characterized |
| *ISM1* | Missense | D181N | Uncharacterized |
| *SLC6A18* | Missense | G33E | Sodium transporter |
| *RNH1* | Missense | D436N | Ribonuclease inhibitor |
| *UBE2R2* | Missense | D232N | Beta-catenin degradation |
| *RHOBTB1* | Missense | V580I | Rho GTPase |
| *MMP15* | Missense | P575Q | Matrix Metallopeptidase |
| *SLC6A13* | Splice donor & intron | Splice site | GABA transporter |

| **Patient 12** | | | Total number of somatic mutations: 114 | |
| --- | --- | --- | --- | --- |
| **Gene** | | **Type of Variant** | **New Mutation** | **Function** |
| *KCMF1* | | Missense | Y82C | E3 Ub Ligase function |
| *SH3PXD2A* | | Missense | T847P | Adapter protein podosome formation |
| *HOXB7* | | Missense | T173M | Homeobox transcription factor |
| *CCDC168* | | Stop gain | S4604* | Uncharacterized |
| *WNK2* | | Missense | R874W | Mitogen-Activated Protein Kinase Kinase Kinase |
| *KRTAP19-8* | | Stop gain | R50* | Keratin associated protein |
| *FAM198A* | | Missense | P42Q | Uncharacterized |
| *KIR3DL2* | | Missense, frame-shift ins & del | N239S | Killer cell immunoglobulin-like receptor |
| *SEC11A* | | Missense | M156I | Peptidase |
| *ANKRD31* | | Stop gain | L276* | Uncharacterized |
| *COPB1* | | Missense | K450Q | Vesicle transport |
| *TEAD4* | | Missense | I242L | Transcription factor, hippo pathway, EMT |
| *SKOR2* | Missense | | F947S | Transcriptional repressor |
| *DENND5B* | | frameshift | AKRTG461VWLW…, | Guanine nucleotide exchange factor |
| *MUC2* | | Missense | S1943R | Member of the mucin family |

| **Patient 14** | | Total number of somatic mutations: 63 | |
| --- | --- | --- | --- |
| **Gene** | **Type of Variant** | **New Mutation** | **Function** |
| *UBE2J1* | Missense & splice region | Y227H | Ubiquitin conjugating enzyme |
| *TET2* | Stop gain | S774* | Methylcytosine Dioxygenase |
| *SLC39A11* | Missense | S153R | Metal ion transporter |
| *SPATA21* | Missense | R227C | Differentiation of haploid spermatids |
| *PRR36* | Missense | P832H | Uncharacterized |
| *GOLGA8K* | Missense | L311V | Uncharacterized |
| *TAAR1* | Missense | K188N | G-Protein coupled receptor |
| *OR2W1* | Missense | I256L | Olfactory receptor |
| *HMCN2* | Missense | G3327S | Calcium ion binding |

| **Patient 13** | | Total number of somatic mutations: 89 | |
| --- | --- | --- | --- |
| **Gene** | **Type of Variant** | **New Mutation** | **Function** |
| *CYP24A1* | Missense | Y145H | Cytochrome p 450 |
| *MRPL46* | Missense | R7W | Mitochondrial ribosomal protein |
| *RIBC2* | Missense | R112C | Uncharacterized |
| *INF2* | Missense | P1163H | (de)polymerization of actin filaments |
| *OR10V1* | Missense | R15C | Olfactory receptor |
| *NKAPL* | Missense | M266I | Transcriptional repressor of Notch signaling |
| *CAMK2G* | Missense | I45T | Calmodulin dependent protein kinase |
| *TANC1* | Missense | D1438N | Not well characterized |

| **Patient 15** | | Total number of somatic mutations: 68 | |
| --- | --- | --- | --- |
| **Gene** | **Type of Variant** | **New Mutation** | **Function** |
| *ATP1A4* | Missense | V287A | Cation transport ATPase |
| *SCLT1* | Missense | T468M | Sodium channel regulator |
| *MCEMP1* | Missense | T159M | Not well characterized |
| *MARCH11* | Missense | R311C | Membrane-bound E3 ubiquitin ligases |
| *CASP5* | Missense | R24H | Aspartate protease (caspase) |
| *ZRANB3* | Missense | M162L | DNA annealing helicase and endonuclease |
| *C5orf30* | Missense | K108E | Putative role in protein trafficking *via* interaction with UNC119 and UNC119B cargo adapters |
| *P2RX7* | Missense | G27V | Ligand gated ion channel, receptor for ATP |
| *LRRC45* | Stop gain | E564* | Centrosome linker required for cohesion |
| *CHRNG* | Missense | C450Y | Muscle-type acetylcholine receptor |

Red, mutations considered to have important cell biological effects.

Supplementary Table 3: Annotated differentially expressed (>2-fold) lncRNAs and miRs in each patient.

| **Patient** | **Differentially expressed lncRNAs** | **Differentially expressed miRs** |
| --- | --- | --- |
| 1 | HOTTIP, SCHLAP1, NEAT1, PCA3, DANCR, MEG3, ARLNC1 | MIR205HG, MIR4300HG, MIR4458HG, MIR646HG, MIR9-3HG, MIR99AHG |
| 2 | SCHLAP1, NEAT1, PCAT1, PCA3, DANCR, MEG3 | MIR1-1HG-AS1, MIR205HG, MIR222HG, MIR23A, MIR27A, MIR5687, MIR5704 |
| 3 | PCAT1, PCA3, DANCR, ARLNC1 | MIR181A1HG, MIR205HG, MIR22HG, MIR27A, MIR3142HG, MIR4458HG, MIR5687, MIR5704, MIR9-3HG |
| 4 | HOTTIP, HOTAIRM1, NKILA, SCHLAP1, PCAT1, PCA3, ARLNC1 | MIR1-1HG-AS1, MIR2052HG, MIR222HG, MIR3142HG, MIR3189, MIR34AHG, MIR3648-2, MIR3687-2, MIR4300HG, MIR4435-2HG, MIR4500HG, MIR4796, MIR5704, MIR578, MIR646HG, MIRLET7BHG |
| 5 | HOTTIP, NKILA | MIR100HG, MIR222HG, MIR3189, MIR4453HG, MIR4458HG, MIR5704 |
| 6 | HOTTIP, NEAT1, ARLNC1 | MIR205HG, MIR3142HG, MIR4429, MIR4435-2HG, MIR561 |
| 7 | HOTTIP, NKILA, SCHLAP1, NEAT1, PCAT1, PCA3, DANCR, MEG3 | MIR1-1HG-AS1, MIR205HG, MIR222HG, MIR22HG, MIR23A, MIR27A, MIR3142HG, MIR5687, MIR5704 |
| 8 | HOTTIP, SCHLAP1, PCAT1, PCA3, DANCR, MEG3 | MIR1-1HG-AS1, MIR205HG, MIR222HG |
| 9 | HOTTIP, NEAT1, PCAT1, PCA3, DANCR, ARLNC1 | MIR100HG, MIR1-1HG-AS1, MIR155HG, MIR193BHG, MIR2052HG, MIR205HG, MIR222HG, MIR3648-2, MIR5704, MIR5706, MIR99AHG, MIRLET7BHG |
| 10 | HOTTIP, SCHLAP1, NEAT1, PCAT1, PCA3, MEG3, ARLNC1 | MIR1-1HG-AS1, MIR222HG, MIR22HG, MIR34AHG, MIR5687, MIR5704 |
| 11 | SCHLAP1, PCAT, ARLNC1 | MIR1251, MIR137HG, MIR22HG, MIR23A, MIR27A, MIR3648-2, MIR578, MIR646HG |
| 12 | HOTTIP, HOTAIRM1, SCHLAP1, NEAT1, PCAT1, PCA3, DANCR, MEG3, ARLNC1 | MIR100HG, MIR1-1HG-AS1, MIR17HG, MIR200CHG, MIR205HG, MIR222HG, MIR25, MIR3142HG, MIR4429, MIR4697HG, MIR503HG, MIR5706, MIR635, MIR646HG, MIR99AHG |
| 13 | NKILA, SCHLAP1, PCAT1, MEG3 | MIR1-1HG-AS1, MIR205HG, MIR22HG, MIR34AHG, MIR4697HG |
| 14 | SCHLAP1, PCAT1, PCA3, MEG3, ARLNC1 | MIR100HG, MIR1-1HG, MIR1-1HG-AS1, MIR1255A, MIR193BHG, MIR205HG, MIR222HG, MIR27B, MIR4697HG, MIR5687, MIR99AHG |
| 15 | NKILA, SCHLAP1, PCAT1, PCA3, MEG3, ARLNC1 | MIR205HG, MIR3189, MIR34AHG, MIR4664, MIR4697HG, MIR5687, MIR646HG, MIR99AHG |

Functionally relevant annotated miRs and lncRNAs were taken from^1-10^. All patient-specific deregulated miRs and lncRNAs can be found in Supplementary File 2. Red, upregulated in tumors; black, down-regulated in tumors relative to the patient-matched normal prostate tissue.

### Supplementary Table 4: Sequencing statistics for RNA-seq and EXOME-seq

| **Normal** | **RNA-SEQ** | | **EXOME-SEQ** | | | **Tumor** | **RNA-SEQ** | | **EXOME-SEQ** | | |
| --- | --- | --- | --- | --- | --- | --- | --- | --- | --- | --- | --- |
|  | Reads | Align % | Reads | Align% | Duplicate reads % |  | Reads | Align% | Reads | Align% | Duplicate reads % |
| 1N_rep1  1N_rep2 | 36603682  38501720 | 92.26  92.64 | 42475200 | 96.07 | 9.12 | 1T_rep1  1T_rep2 | 49485994  52252504 | 91.34  91.78 | 55706200 | 96.12 | 9.12 |
| 2N_rep1  2N_rep2 | 35578814  63894243 | 90.50  91.60 | 56298000 | 96.51 | 11.09 | 2T_rep1  2T_rep2 | 34340289  53856558 | 86.04  91.50 | 44136000 | 96.09 | 10.30 |
| 3N_rep1  3N_rep2 | 33037647  38061308 | 88.49  87.49 | 45167400 | 95.96 | 11.62 | 3T_rep1  3T_rep2 | 39206644  44685255 | 83.38  87.41 | 45636400 | 95.90 | 9.85 |
| 4N_rep1  4N_rep2 | 39635442  54890556 | 87.12  92.23 | 45418800 | 96.31 | 8.21 | 4T_rep1  4T_rep2 | 47273179  53502497 | 84.77  91.40 | 49255300 | 96.06 | 8.41 |
| 5N_rep1  5N_rep2 | 40283379  46638779 | 91.73  90.20 | 49694100 | 96.43 | 7.78 | 5T_rep1  5T_rep2 | 64602422  47515766 | 94.25  91.17 | 43400000 | 96.26 | 13.96 |
| 6N_rep1  6N_rep2 | 46031895  55861482 | 86.04  91.92 | 34157000 | 96.06 | 9.98 | 6T_rep1  6T_rep2 | 33455857  76033143 | 87.00  92.16 | 45420600 | 96.29 | 8.19 |
| 7N_rep1  7N_rep2 | 31882647  42675237 | 87.96  87.76 | 43500400 | 96.04 | 11.28 | 7T_rep1  7T_rep2 | 47721526  38438256 | 86.91  93.03 | 47714500 | 96.02 | 11.74 |
| 8N_rep1  8N_rep2 | 35333114  80370444 | 83.44  94.85 | 54276700 | 96.33 | 9.09 | 8T_rep1  8T_rep2 | 74598149  44121100 | 88.50  91.54 | 48035100 | 96.09 | 14.18 |
| 9N_rep1  9N_rep2 | 80833193  41562421 | 88.13  94.28 | 46282300 | 95.93 | 10.72 | 9T_rep1  9T_rep2 | 39677083  38703772 | 90.05  93.42 | 48773100 | 96.04 | 7.44 |
| 10N_rep1  10N_rep2 | 53431330  51151259 | 85.30  91.93 | 53645600 | 96.29 | 8.11 | 10T_rep1  10T_rep2 | 42928847  54227448 | 86.65  91.26 | 55294500 | 96.26 | 8.08 |
| 11N_rep1  11N_rep2 | 42821733  56544524 | 83.82  92.71 | 51965800 | 96.32 | 9.47 | 11T_rep1  11T_rep2 | 48657860  53899463 | 82.43  92.03 | 57373400 | 96.42 | 11.73 |
| 12N_rep1  12N_rep2 | 37996964  40528066 | 91.08  91.61 | 55279200 | 96.12 | 16.20 | 12T_rep1  12T_rep2 | 44233768  47182837 | 91.96  92.44 | 52977500 | 96.37 | 8.51 |
| 13N_rep1  13N_rep2 | 48075036  44318278 | 87.48  93.21 | 49494300 | 95.97 | 5.36 | 13T_rep1  13T_rep2 | 51068154  48949742 | 87.50  91.89 | 60609000 | 96.16 | 11.58 |
| 14N_rep1  14N_rep2 | 36167783  45548318 | 86.15  90.53 | 46894500 | 96.11 | 9.67 | 14T_rep1  14T_rep2 | 45615157  35440307 | 85.63  89.91 | 56130300 | 96.08 | 14.89 |
| 15N_rep1  15N_rep2 | 46655594  43936010 | 88.21  90.63 | 53319000 | 96.06 | 12.62 | 15T_rep1  15T_rep2 | 43724856  66277516 | 85.14  91.99 | 49329900 | 96.03 | 11.78 |

Strand specific paired-end RNA sequencing (Illumina Hiseq 2500, 125/150bp) of Ribo-depleted total RNA from biological duplicates (adjacent tissues sections). Quality of sequencing was assessed using FastQC tool. Exome sequencing was performed using Exome capture kit (Agilent) and subsequently sequenced on Hiseq2500 for 125 bp paired-end.

### Supplementary Table 5: PCR primer sequences

| Gene name | Forward Primer 5’ 🡪 3’ | Reverse Primer 5’ 🡪 3’ |
| --- | --- | --- |
| MGA | CGAAAGAGCCGGGGTGAGAA | TGCCTCCATGTAATCTCTCCCT |
| CFTR | TCCTAACTGAGACCTTACACCGT | AGGCTCATCAGAATCCTCTTCGA |
| MST1R | TCACCCAGTGAGAAGTTGG | ACCTATTCAATGGGCTGTTG |
| RASSF8 | GACAGAGAAGCTTCAATCCA | GAAGTTGATCCTTCAGCTG |

Primers used for validation of Exome-seq data using sanger sequencing.

### Access to WES Analysis Pipeline and custom scripts

The WES Analysis Pipeline script is available at the following link for download:

<https://drive.google.com/open?id=1P9iORvMSAAiVahk0XA0vhOkRCinKOjSM>

For generating STRING network for each patient using a list of genes: <https://drive.google.com/open?id=1G4RRWMddV80ZfyecLHP-sYJpfQUx3Ma4>

For generating per patient gene information whether it's DE/mutated/both: <https://drive.google.com/open?id=15yDE8hUUS9CkFhRCPy8XgtxhnTSKeMqf>

For adding gene information of each patient to STRING network: <https://drive.google.com/open?id=1Mw79BGhW4RnjOpRT4eh7MAEqGO5GhQRd>

For removing duplicate lines from CellNet network: <https://drive.google.com/open?id=1L0387lNi3hw5L9GYfTVlwgYMH0xm-tyB>

### URLs

qcGenomics: http://ngs-qc.org/qcgenomics/;

TETRAMER: <http://www.ngs-qc.org/tetramer/>;

CellNet; <http://cahanlab.org/resources.html> ; <https://github.com/pcahan1/CellNet>

STRING; <https://string-db.org/> ;

CYTOSCAPE: <https://cytoscape.org/>;

MUTECT2:<https://software.broadinstitute.org/gatk/documentation/tooldocs/3.80/org_broadinstitute_gatk_tools_walkers_cancer_m2_MuTect2.php> ;

PANTHER: <http://pantherdb.org/>;

annoPeakR: https://apps.medgen.iupui.edu/rsc/content/19/;

SAMtools 1.6: https://sourceforge.net/projects/samtools/files/samtools/1.6/;

GATK: https://software.broadinstitute.org/gatk/;

Picard 2.14: <http://broadinstitute.github.io/picard/>;

FastQC: <http://www.bioinformatics.babraham.ac.uk/projects/fastqc>;

Hisat2: <https://ccb.jhu.edu/software/hisat2/index.shtml>;

DEseq2 (1.20.0): <https://bioconductor.org/packages/release/bioc/html/DESeq2.html>;

SnapGene Viewer: <https://www.snapgene.com/snapgene-viewer/>;

SummarizedExperiment:<https://www.bioconductor.org/packages/devel/bioc/vignettes/SummarizedExperiment/inst/doc/SummarizedExperiment.html>;

dbSNP: https://www.ncbi.nlm.nih.gov/snp/

### References

1.	Arun, G., Diermeier, S.D. & Spector, D.L. Therapeutic Targeting of Long Non-Coding RNAs in Cancer. *Trends Mol Med* **24**, 257-277 (2018).

2.	Schmitt, A.M. & Chang, H.Y. Long Noncoding RNAs in Cancer Pathways. *Cancer Cell* **29**, 452-463 (2016).

3.	Campbell, J.D. *et al.* Genomic, Pathway Network, and Immunologic Features Distinguishing Squamous Carcinomas. *Cell Rep* **23**, 194-212 e6 (2018).

4.	Di Agostino, S. *et al.* Long Non-coding MIR205HG Depletes Hsa-miR-590-3p Leading to Unrestrained Proliferation in Head and Neck Squamous Cell Carcinoma. *Theranostics* **8**, 1850-1868 (2018).

5.	Larzabal, L. *et al.* TMPRSS4 regulates levels of integrin alpha5 in NSCLC through miR-205 activity to promote metastasis. *Br J Cancer* **110**, 764-74 (2014).

6.	Li, X. *et al.* Dissecting LncRNA Roles in Renal Cell Carcinoma Metastasis and Characterizing Genomic Heterogeneity by Single-Cell RNA-seq. *Mol Cancer Res* **16**, 1879-1888 (2018).

7.	Pipan, V., Zorc, M. & Kunej, T. MicroRNA Polymorphisms in Cancer: A Literature Analysis. *Cancers (Basel)* **7**, 1806-14 (2015).

8.	Sarkozy, M., Kahan, Z. & Csont, T. A myriad of roles of miR-25 in health and disease. *Oncotarget* **9**, 21580-21612 (2018).

9.	Wan, X. *et al.* Androgen-induced miR-27A acted as a tumor suppressor by targeting MAP2K4 and mediated prostate cancer progression. *Int J Biochem Cell Biol* **79**, 249-260 (2016).

10.	Wang, C. *et al.* MiR-25 promotes hepatocellular carcinoma cell growth, migration and invasion by inhibiting RhoGDI1. *Oncotarget* **6**, 36231-44 (2015).
